# Cooperative [2Fe-2S] cluster-binding regulates the functional transitions of the *Aspergillus fumigatus* iron regulator HapX for adaptation to iron starvation, sufficiency and excess

**DOI:** 10.1101/2024.11.27.625597

**Authors:** Simon Oberegger, Matthias Misslinger, Hubertus Haas

## Abstract

Accurate sensing of cellular iron levels is vital, as this metal is essential but toxic in excess. The iron-sensing transcription factor HapX is crucial for virulence of *Aspergillus fumigatus,* the predominant human mold pathogen. Its absence impairs growth under iron limitation and excess, but not under moderate iron availability, suggesting that HapX switches between three states to adapt to varying iron availability.

This study suggests that the HapX state transitions are regulated by the different propensities of four phylogenetically conserved cysteine-rich regions (CRRs) to coordinate [2Fe-2S] clusters resulting in cumulative occupancies that depend on iron availability. In the iron starvation state, CRR-B and -C lack [2Fe-2S] clusters, the iron sufficiency/”neutral” state features clusters in CRR-B and/or -C and the iron excess state has clusters in all CRR-A, B, and -C, while CRR-D plays a minor role. Combinatorial mutation of CRR-B and -C blocked growth by locking HapX in the iron starvation state, leading to uncontrolled iron uptake, iron accumulation, repression of iron-consuming pathways and impaired iron detoxification. Loss of the C-terminal 27 amino acid region of HapX, which is crucial for the iron starvation state and was found to contain a degron, rescued the severe growth defect. Noteworthy, the - Fe state of HapX induced several gene clusters encoding secondary metabolites.

## INTRODUCTION

Iron is an indispensable trace element which is required by all eukaryotic and virtually all prokaryotic species ^1^. The ability of iron to switch between two oxidation states, ferrous iron (Fe^2+^) and ferric iron (Fe^3+^), has made it an essential cofactor in redox biochemistry as heme, siroheme, iron-sulfur clusters, and mono- or dinuclear iron centers ^2–5^. Iron-dependent pathways include respiration, the TCA cycle, oxidative stress detoxification, P450 enzymes, DNA repair and replication as well as the biosynthesis of amino acids, nucleotides and sterols. Moreover, iron is also critically involved in less conserved pathways such as secondary metabolism ^4^. Although iron is essential, excess iron leads to the formation of highly reactive hydroxyl radicals through Haber-Weiss-Fenton chemistry causing oxidative cell damage ^6,7^. Therefore, sophisticated mechanisms are required to balance iron uptake, consumption, and storage to avoid both deficiency and toxicity of this metal, i.e. organisms must be able to discriminate cellular iron limitation (-Fe), iron sufficiency (+Fe) and iron excess (hFe) for respective adaptation.

*Aspergillus fumigatus* is one of the most common opportunistic fungal pathogens causing life-threatening invasive infectious diseases, particularly in immunocompromised patients, known as aspergillosis ^8,9^. Iron homeostasis has been shown to be key to the virulence of this mold in various infection models ^4^. In *A. fumigatus* iron homeostasis is mainly regulated by two iron-responsive transcription factors, termed HapX and SreA, which are interconnected in a negative transcriptional feedback loop ^4^. In -Fe, HapX represses iron-consuming pathways, including heme biosynthesis, respiration and vacuolar iron detoxification, while inducing high-affinity iron uptake through siderophore-mediated and reductive iron assimilation ^10–12^. hFe turns HapX into an activator of iron-consuming pathways and vacuolar iron deposition by the transporter CccA ^13,14^. During sufficient iron supply, SreA represses expression of HapX and high-affinity iron acquisition by siderophore-mediated and reductive iron assimilation ^15,16^. Consequently, loss of HapX impairs growth in -Fe and hFe but not +Fe conditions, whereas loss of SreA impairs growth in +Fe and especially in hFe but not in -Fe conditions 10,14,15,17.

SreA, a GATA-type zinc finger transcription factor, contains a single cysteine-rich region (CRR) for iron sensing ^15,18–20^. HapX shows several, evolutionary highly conserved characteristics: (i) a Hap4-like domain mediating direct interaction with the CCAAT-binding complex for cooperative DNA binding ^12,21,22^, (ii) the bZIP-type DNA-binding domain, (iii) a coiled-coil domain for dimerization, (iv) four CRRs, termed CRR-A to CRR-D that are involved in iron sensing, and (v) the C-terminal 27 amino acid region required for the role of HapX in -Fe but not hFe ^4,14,21^. Each CRR of SreA and HapX contains four cysteine residues, which differ in spacing and amino acid composition but show high phylogenetic conservation ^4,23^. Several lines of evidence indicate that regulation of iron homeostasis via SreA and HapX depends on [2Fe-2S] cluster sensing by the CRRs: (i) inactivation of mitochondrial [2Fe-2S], but not cytosolic [4Fe-4S] cluster biogenesis, impaired iron sensing in *A. fumigatus* ^24^; (ii) mutational analysis revealed that mainly CRR-B, but to a lesser extent also CRR-A and even less CRR-C, are required for HapX’s role in hFe but not -Fe, while mutation of CRR-D was phenotypically inconspicuous ^14^; (iii) GrxD, which belongs to the [2Fe-2S] cluster-transferring monothiol glutaredoxins ^25^, physically interacts with both HapX and SreA and appears to mediate the removal of [2Fe-2S] clusters from both HapX and SreA to switch HapX from the “iron” to the “-Fe” state and to inactivate SreA ^17^; (iv) the C-terminal domain of HapX (amino acids 161–491) containing the four CRRs, recombinantly expressed in *Escherichia coli*, exhibits a UV-vis spectrum indicative of [2Fe-2S] clusters ^17^; and (v) peptide mimics of the CRRs of HapX and SreA have the ability to coordinate [2Fe-2S] clusters *in vitro* with differing propensities, whereby the [2Fe-2S] cluster coordinated by HapX CRR-B showed an outstanding high stability ^23^. HapX and SreA are highly conserved in the fungal kingdom but not in *Saccharomycotina*, which employ different iron-sensing transcription factors ^17^. Remarkably, the GrxD homologous monothiol glutaredoxins Grx3 and Grx4 are required for adaptation to iron sufficiency in *Saccharomyces cerevisiae* in contrast to the situation in *A. fumigatus*, where GrxD is important for -Fe adaptation ^17,26,27^. These data highlight significant differences between iron homeostatic systems, even within fungal species.

Genetic inactivation of HapX causes growth and metabolic defects under -Fe and hFe but not +Fe ^10,14^, indicating that this regulator can switch between three distinct states, mediating adaptation to -Fe, +Fe (neutral/inactive) and hFe. Similar to siderophore biosynthesis and uptake ^28–31^, HapX is crucial for *A. fumigatus* virulence in a murine model of aspergillosis, highlighting -Fe in the host niche and the importance of pathogen adaptation ^10^. The latter demonstrates that switching HapX into its -Fe state is important for adaptation to the mammalian host niche. However, the molecular determinants of the HapX -Fe state have remained elusive. In this study, we elucidated the molecular mechanism for the HapX transitions between its three distinct functional states for adaptation to -Fe, +Fe (neutral/inactive), and hFe.

## RESULTS

### Combinatorial mutation of HapX CRR-A, CRR-B, CRR-C and CRR-D blocks growth

Previously, mutational analysis in HapX revealed that mainly CRR-B, but to a lesser extent also CRR-A and even less CRR-C, are required for HapX’s role in hFe but not -Fe, while mutation of CRR-D was phenotypically inconspicuous ^14^. Important to note, individual replacement of different cysteine residues of the same CRR by alanine had the same phenotypical consequences, indicating that the four cysteine residues in a CRR function in concert and that replacement of a single cysteine residue is sufficient to inactivate the function of a CRR ^14^. Attempts to generate a HapX allele expressed under its native promoter and lacking all four CRRs were unsuccessful, suggesting that this might be lethal. For further functional analysis of HapX protein domains, generated *hapX* alleles were therefore expressed under control of the xylose-inducible *xylP* promoter (*PxylP*), which allows tunable expression levels, including silencing and overexpression ^32,33^, with single copy integration at the native *hapX* locus. For phenotyping, fungal strains were grown on solid media reflecting -Fe, +Fe and hFe without xylose for repression and 0.1% xylose supplementation for expression of *PxylP*-driven *hapX* alleles (Figure 1). Compared to +Fe, *A. fumigatus* wildtype (*wt*) displayed reduced conidiation (reduced green coloration of the fungal colony) in -Fe and reduced radial growth in hFe, largely independent of xylose supplementation. Compared to *wt*, lack of HapX (*ΔhapX*) decreased conidiation in -Fe and blocked growth in hFe but showed *wt*-like growth in +Fe, independent of xylose supplementation, as previously reported ^14^. Without xylose supplementation, *PxylP*-driven native *hapX* displayed a *ΔhapX* growth phenotype, apart from slight growth in hFe most likely due to slight leakiness of *PxylP* ^33^. With xylose induction, *PxylP*-driven native *hapX* displayed a largely *wt*-like growth pattern (apart from mildly decreased growth in hFe), demonstrating the suitability of *PxylP* for studying functionality of *hapX* alleles.

**Figure 1:**
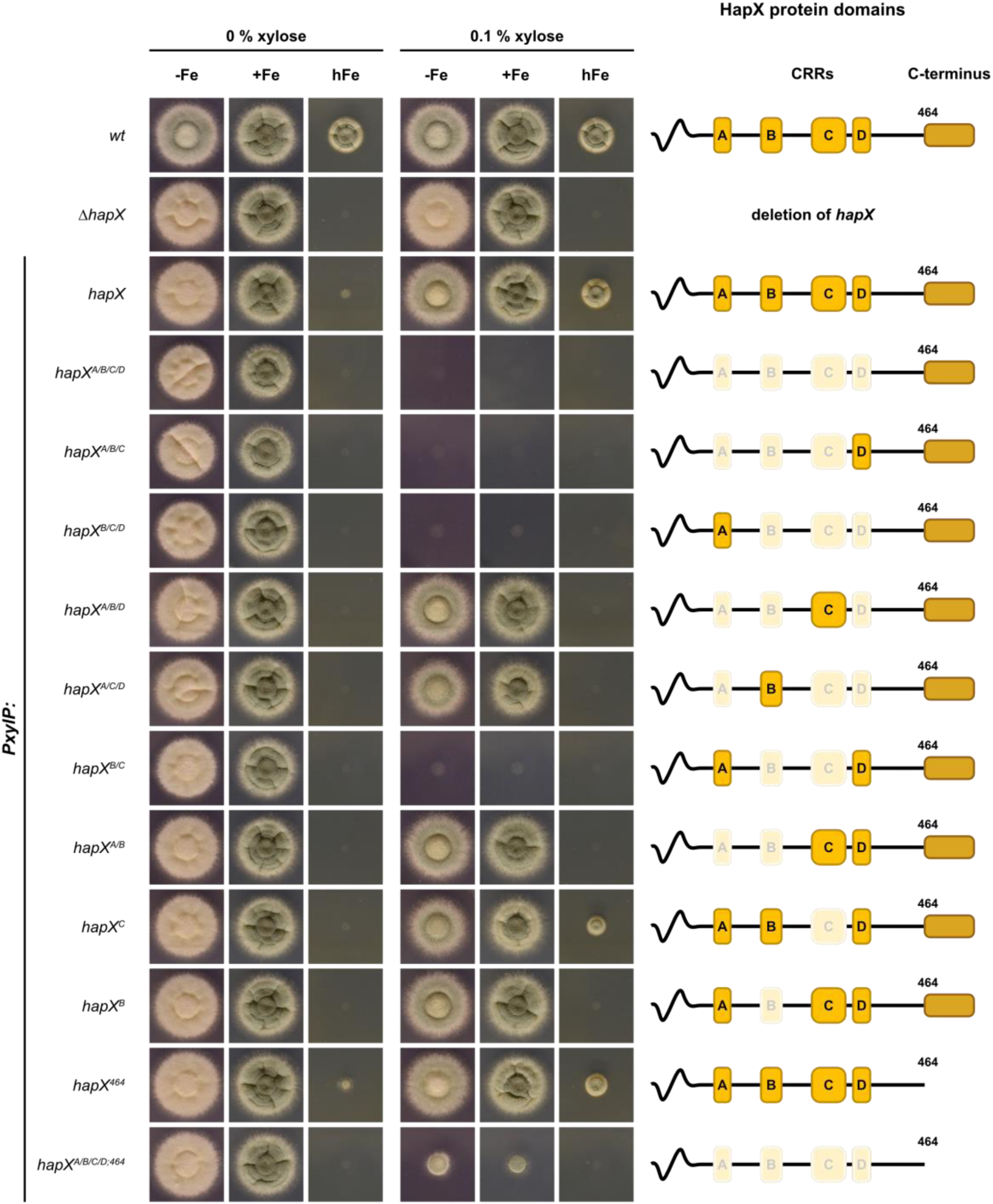
Combinatorial mutation of CRR-B and -C (strains *hapX^A/B/C/D^*, *hapX^A/B/C^*, *hapX^B/C/D^*, *hapX^B/C^*), but not their individual mutation (*hapX^B^*, *hapX^C^*), blocks growth independent of iron availability (-Fe / +Fe / hFe). For every fungal strain, 10^4^ conidia were spot-inoculated on minimal medium plates reflecting iron starvation (-Fe; 200 µM BPS), iron sufficiency (+Fe; 30 µM FeSO_4_) or excess iron (hFe; 10 mM FeSO_4_). Expression of *PxylP*-driven *hapX* alleles was tuned by supplementation with xylose: 0% for repression and 0.1% for moderate expression. Plates were incubated at 37 °C for 48 h. The CRRs and the C-terminus of the respective HapX alleles are schematically shown on the ride side; functional inactivation of a CRRs is indicated by white shading.

As a first step in the functional analysis of the HapX CRRs, we simultaneously replaced one cysteine of each CRR by an alanine residue (C203A, C277A, C353A, C380A) to block [2Fe-2S] cluster binding by all four CRRs, resulting in strain *hapX^A/B/C/D^*. Under non-inducing conditions (0% xylose), this strain showed a growth pattern similar to that of *ΔhapX* (Figure 1). Since the latter growth behavior was observed for all generated mutant strains without xylose supplementation (Figure 1), it is not discussed further for the other generated strains. Remarkably, xylose-induced expression of the *hapX^A/B/C/D^* allele blocked growth under all conditions tested (Figure 1). Importantly, individual mutation of the four CRRs of HapX expressed under the endogenous promoter was previously shown to negatively affect growth only in hFe, whereby this was observed mainly for CRR-B, less for CRR-A, only mildly for CRR-C and not at all for CRR-D ^14^. Similarly, under *PxylP* control, individual mutation of CRR-B (strain *hapX^B^*) blocked and that of CRR-C (strain *hapX^C^*) mildly decreased growth exclusively in hFe compared to the native *hapX* allele with xylose induction (Figure 1).

To investigate the reason for the growth blocking effect of *hapX^A/B/C/D^*allele expression, we compared the expression of HapX target genes in strains expressing the *PxylP*-driven native *hapX* and *hapX^A/B/C/D^* alleles by Northern blot analysis. Therefore, *hapX* and *hapX^A/B/C/D^* strains were grown in +Fe liquid cultures without xylose supplementation for 17 h to circumvent the growth inhibitory effect of combinatorial mutation of all four CRRs, taking advantage of the phenotypically inconspicuous *ΔhapX* phenotype in +Fe (Figure 1), followed by incubation for 1 h in the presence of varying amounts of xylose to induce expression of the *hapX* alleles. Additionally, the *wt* was grown for 18 h in -Fe and +Fe liquid cultures as control. Harvested mycelia were subjected to Northern blot analysis of iron-regulated genes. In line with previously reported data ^10,14^, *hapX* transcript levels were strongly upregulated in -Fe compared to +Fe in *wt* (Figure 2A). In contrast, strains with *PxylP*-driven *hapX* showed *hapX* expression that increased with xylose supplementation from 0.05 to 1.0% xylose, whereas *hapX* transcripts were not detected without (0%) or low (0.01%) xylose supplementation (Figure 2B). In contrast to the *hap*X strain, with xylose induction ≥0.05%, the *hapX^A/B/C/D^* strain showed an expression pattern similar to that of the *wt* grown in -Fe (Figure 2B), i.e. induction of -Fe-induced *mirB*, which mediates siderophore uptake ^34^ and repression of -Fe-repressed heme biosynthetic *hemA* and iron regulatory *sreA* (Figure 2B). This expression pattern of strain *hapX^A/B/C/D^* is consistent with the previously reported -Fe response in *A. fumigatus* ^10,14,15,17^. These data strongly suggest that the simultaneous mutational disruption of all four HapX CRRs renders HapX iron-blind, regardless of the prevailing iron availability, as shown here for +Fe. As shown previously ^15,18^, *sreA* expression resulted in two different sized transcripts due to alternative transcription initiation. Notably, expression of *hapX^A/B/C/D^* mainly affected the larger *sreA* transcript (Figure 2B), revealing differential transcriptional regulation of the two *sreA* transcripts by HapX. Remarkably, high expression (1.0% xylose) of the *wt hapX* allele resulted in weak induction of *mirB* (Figure 2B).

**Figure 2:**
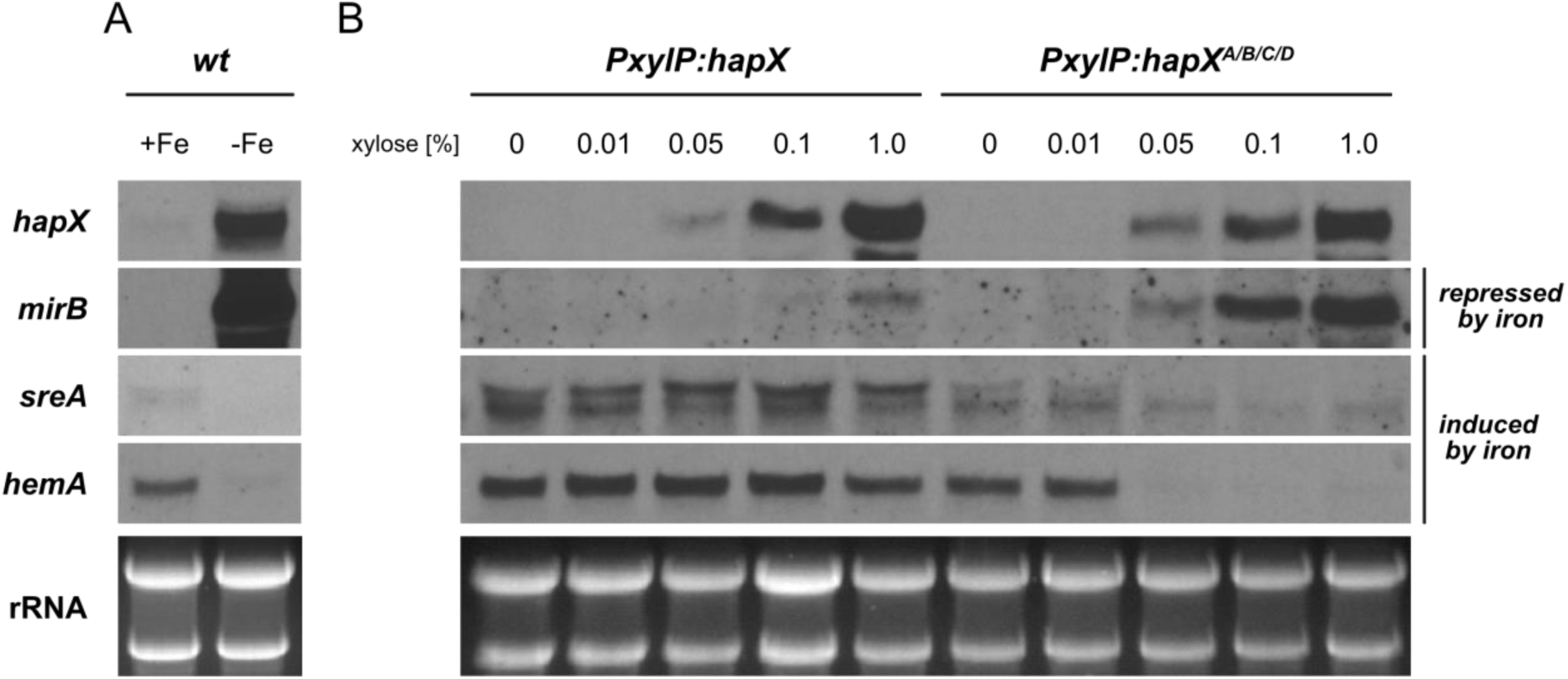
Combinatorial mutation of all four HapX CRRs (*hapX^A/B/C/D^*) results in an iron starvation response in +Fe. Total RNA was isolated from liquid cultures grown either in +Fe or -Fe at 37 °C. The *wt* was grown for 18 h (A), strains expressing *PxylP*-driven *hapX* and *hapX^A/B/C/D^*were grown in +Fe without xylose for 17 h followed by 1 h incubation with different xylose concentrations for induction of *hapX* expression (B). Northern blot analyzed genes: *hapX*, iron limitation-induced *mirB* as well as iron limitation-repressed *sreA* and *hemA*. Ethidium bromide-stained rRNA is shown for loading and quality contol.

### Combinatorial mutation of the CRRs locks HapX in the -Fe status causing cellular stress and affecting secondary metabolism

The Northern blot analysis of selected genes shown above (Figure 2) cumulatively suggested that combinatorial mutation of the four CRRs locks HapX in the -Fe state. To characterize the genome-wide effect of combinatorial mutation of the four CRRs, we employed high-throughput RNA sequencing for transcriptome profiling. Therefore, biological triplicates of strains encoding *PxylP*-driven *hapX* and the *hapX^A/B/C/D^* were grown in liquid culture in +Fe for 17 h to allow growth under non-inducing conditions followed by an incubation for 1 h in the presence of 0.1% xylose for induction of the *hapX* alleles. On average, we obtained 23,476,866 reads per individual sample. Genome alignment revealed an average unique mapping percentage of 82.5% to the *A. fumigatus* Af293 reference genome, identifying the expression of 9,859 genes, of which 1,406 were found to be upregulated and 1,175 were downregulated ≥ 2-fold in strain *hapX^A/B/C/D^* compared to *hapX* with an adjusted p-value < 0.05 (Figure 3). The transcriptome data are shown in Supplementary Table S1A; principal component and volcano plot analyses of the transcriptomic comparison are shown in Supplementary Figure S1.

**Figure 3:**
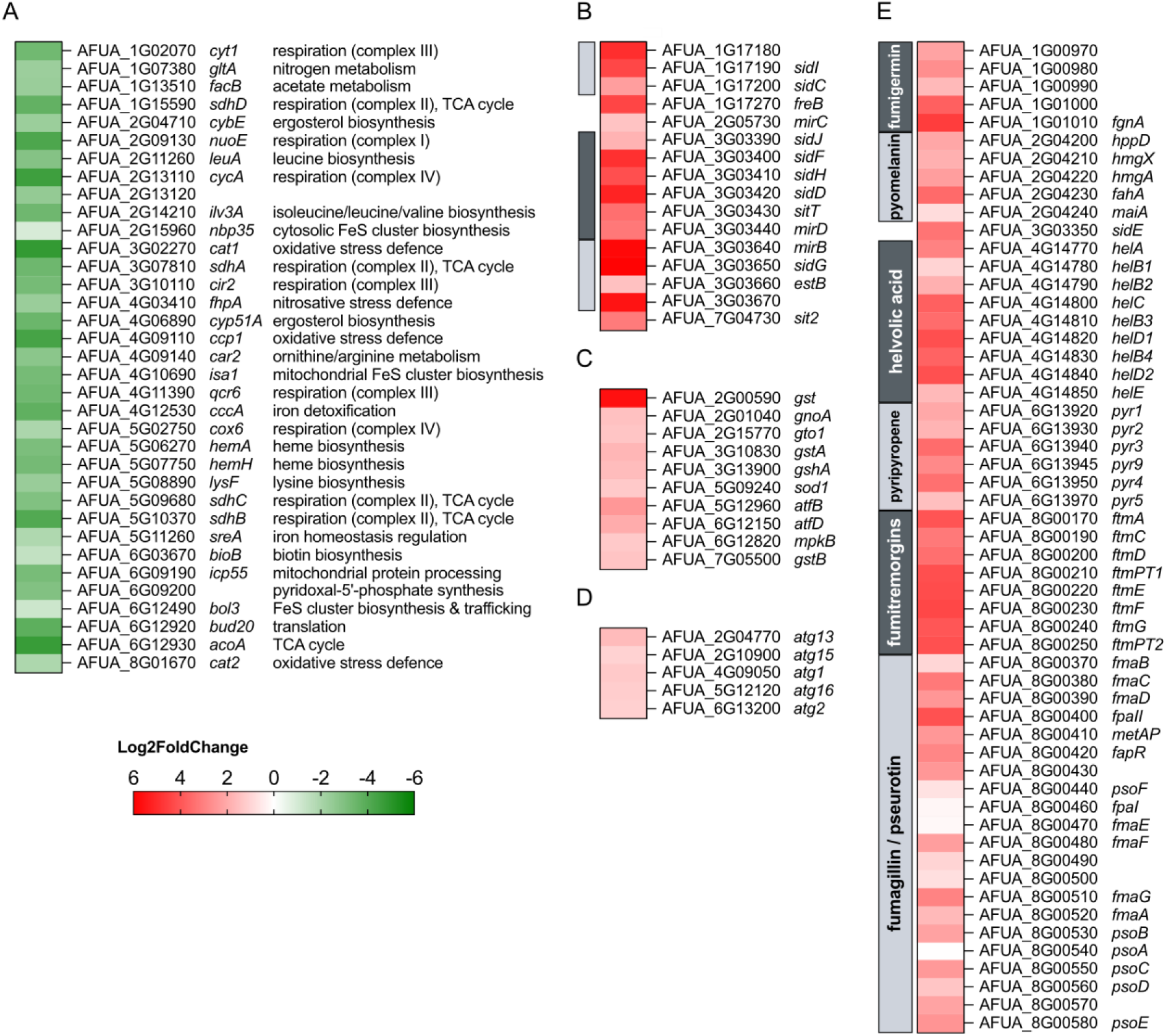
Heatmap comparison of selected differentially expressed genes in *hapX^A/B/C/D^* compared to *hapX* in +Fe. Genes that have previously been shown to be directly downregulated by HapX in -Fe (A); genes involved in siderophore metabolism or localized in siderophore metabolic gene clusters that have previously been shown to be upregulated in -Fe (B); genes involved in stress response (C); genes involved in autophagy (D); and genes involved in secondary metabolism (E). Only genes downregulated with a log2 fold change ≤ -1 or upregulated with ≥ 1 are shown, with the exception of (E): here, all genes of respective secondary metabolite encoding gene clusters are shown. Fumarylalanine is synthesized by SidE, most likely without the need for other enzymes. Gene clusters are highlighted by grey shadded boxes on the left side of the heat maps. The respective transcriptome data subsets are shown in Supplementary Table S1B.

A previous study identified 33 genes that are directly repressed by the CBC:HapX complex in - Fe ^12^. These genes encode iron-linked steps in various pathways including respiration, TCA cycle, CccA-mediated vacuolar iron deposition, defense against oxidative and nitrosative stress by catalase and flavohemoglobin, SreA, as well as biosynthesis of heme, iron-sulfur clusters and amino acids. All of these genes were downregulated in *hapX^A/B/C/D^*compared to *hapX* (Figure 3A). Moreover, the majority of genes involved in siderophore-mediated iron assimilation or localized in encoding gene clusters, that have been previously shown to be repressed by SreA ^15,17^, were upregulated in *hapX^A/B/C/D^*compared to *hapX* (Figure 3B). Taken together, these results demonstrate that combinatorial mutation of the four CRRs locks HapX in the -Fe state leading to a transcriptional -Fe response in +Fe conditions. Therefore, the most likely reason for the severe effect of combinatorial mutation of the four CRRs described above is caused by uncontrolled iron acquisition combined with repression of iron-consuming pathways and CccA-mediated vacuolar iron detoxification.

Iron overload is expected to cause cellular stress, particularly oxidative and nitrosative stress due to the formation of highly reactive oxygen and nitrogen species, also known as Haber-Weiss-Fenton and Fenton-like chemistry ^6,7,35,36^. In line, several genes encoding stress-responsive regulatory or structural proteins were found to be upregulated in strain *hapX^A/B/C/D^* (Figure 3C), including the mitogen-activated protein kinase MpkB ^37^, the basic leucine zipper (bZIP) transcription factors AtfB and AtfD ^38^, glutathione biosynthetic GshA ^24^, several glutathione transferases (Gto1, GstA, GstB; ^39^, Cu/Zn-superoxide dismutase SodA ^40^ and nitrosative stress-detoxifying S-nitrosoglutathione reductase GnoA ^41^. Noteworthy, the genes encoding oxidative stress-detoxifying mycelial catalases Cat1 and Cat2 ^42^ as well as nitrosative stress-detoxifying flavohemoglobin FhpA ^41^ were significantly downregulated in strain *hapX^A/B/C/D^* (Figure 3A). However, these genes have previously been shown to be directly repressed by HapX during -Fe ^12,14^. Consequently, the activation of the -Fe state of HapX in +Fe appears to result not only in uncontrolled iron uptake combined with repression of iron consumption and detoxification but most likely also impairs defense against oxidative and nitrosative stress.

Remarkably, the majority of the downregulated genes in strain *hapX^A/B/C/D^* are related to ribosome biogenesis and translation (Supplementary Figure S2 and Supplementary Table S1C), while five genes involved in autophagy were upregulated (Figure 3D), a pattern consistent with repression of TOR complex 1 (TORC1; target of rapamycin) ^43,44^.

In addition, biosynthetic gene clusters for the secondary metabolites fumigermin ^45^, pyomelanin/tyrosine degradation ^46^, fumarylalanine ^47^, helvolic acid ^48^, the meroterpenoid pyripyropene ^49^, fumitremorgins ^50^, and fumagillin/pseurotin ^51,52^ were upregulated in strain *hapX^A/B/C/D^*.

Notably, induction of fumarylalanine and pyomelanin/tyrosine degradation has previously been linked to cellular stress ^46,47^. In contrast, expression of gene clusters ^53^ for biosynthesis of fumigaclavines (*fga*), hexadehydroastechrome (*has*), isocyanides fumivaline/fumicicolin (*crm*), endocrocin (*enc*), trypacidin (*tpc*), xanthozillin (*xan*), fumisoquin (*fmp/fsq*), gliotoxin (*gli*), neosartoricin/fumicycline (*nsc*), trypacidin (*tpc*), fumihopaside (*afum*), satorypyrone (spy) was largely unaffected in strain *hapX^A/B/C/D^* compared to *hapX* (Supplementary Table S1A).

### Combinatorial mutation of HapX CRR-B and CRR-C blocks growth and activates the -Fe state

To elucidate the combinatorial mutation of which of the four CRRs is responsible for the growth inhibitory effect, i.e. which CRRs are involved in locking HapX in the -Fe state, we generated mutant strains expressing *hapX* alleles carrying all combinatorial mutations of three CRRs, termed *hapX^A/B/C^*, *hapX^B/C/D^*, *hapX^A/B/D^*, and *hapX^A/C/D^*. Phenotyping revealed that only *hapX^A/B/C^* and *hapX^B/C/D^*strains phenocopied the *hapX^A/B/C/D^* strain upon xylose induction (Figure 1), suggesting that the combinatorial mutation of CRR-B and -C, which is common to these two alleles, is most likely responsible for the growth defect. Indeed, combinatorial mutation of only CRR-B and -C (strain *hapX^B/C^*) blocked growth upon xylose induction, regardless of the prevailing iron availability (Figure 1). In contrast, individual mutation of CRR-B (strain *hapX^B^*) or CRR-C (strain *hapX^C^*) or combinatorial mutation of CRR-A and -B (strain *hapX^A/B^*) impaired growth only in hFe (Figure 1). Taken together, these results demonstrate that combinatorial mutation of CRR-B and -C is sufficient to block growth, regardless of the prevailing iron availability. As previously shown in strains expressing h*apX* under its native promoter ^14^, individual mutation of CRR-B (strains *hapX^B^, hapX^A/B^*, *hapX^A/B/D^*) blocked and mutation of CRR-C (strain *hapX^C^*) reduced growth exclusively in hFe (Figure 1). Moreover, combinatorial mutation of CRR-A, -C and -D (strain *hapX^A/C/D^*) blocked growth in hFe (Figure 1). These results highlight the importance of CRR-A, -B and -C in mediating iron resistance in *A. fumigatus,* as previously demonstrated with individual mutation of CRRs ^14^. Mutation of CRR-D was phenotypically inconspicuous in all combinations tested (Figure 1).

All generated strains expressing *hapX* alleles were subjected to Northern blot analysis in +Fe with xylose induction for 1 h (Figure 4). In contrast to *ΔhapX*, all strains expressing *PxylP*-driven *hapX* alleles showed largely similar *hapX* induction (Figure 4A), whereby *hapX* transcript levels were slightly higher in strains *hapX^A/B/C/D^*, *hapX^A/B/C^*, *hapX^B/C/D^* compared to the other strains. Similar to the strain expressing *hapX* with combinatorial mutation of CRR-A, -B, -C and -D (*hapX^A/B/C/D^*), all other strains with combinatorial mutation of CRR-B and -C (strains *hapX^A/B/C^*, *hapX^B/C/D^*and *hapX^B/C^*), in contrast to the remaining strains (*ΔhapX, hapX, hapX^A/B/D^, hapX^A/C/D^, hapX^A/B^, hapX^C^,* and *hapX^B^*), showed a transcriptional -Fe response in +Fe, i.e. higher transcript levels of *mirB* and lower transcript levels of *hemA* and the larger *sreA* transcript (Figure 4A), similar to *wt* in -Fe (Figure 4B). Again, mutation of CRR-D did not affect the transcriptional pattern in any combination. Taken together, these data underscore that the combinatorial mutation of CRR-B and -C blocks growth by causing "iron blindness" of HapX. In turn, these data suggest that CRR-B and -C do not coordinate [2Fe-2S] clusters in -Fe and that [2Fe-2S] cluster occupancy of CRR-B and/or -C switches HapX out of the iron starvation state.

**Figure 4:**
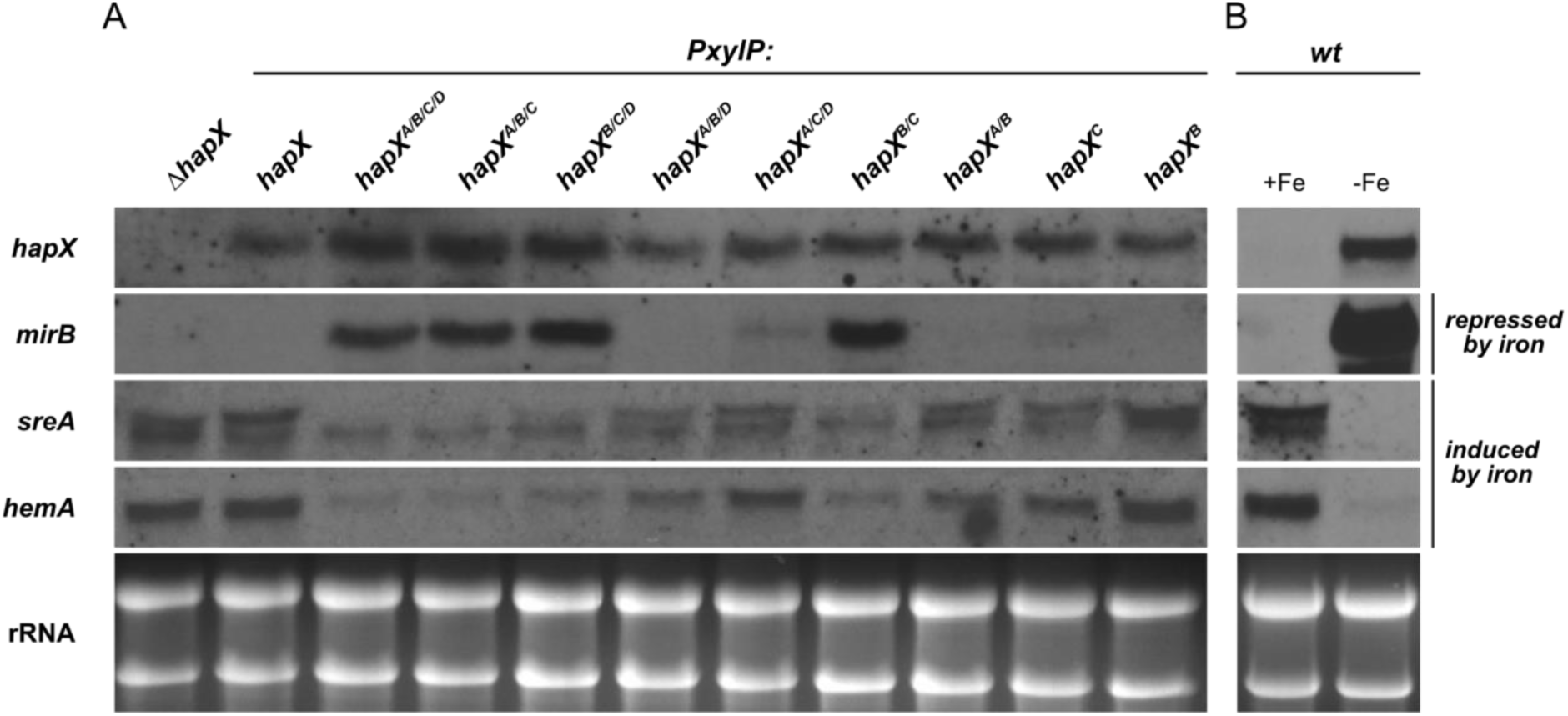
Combinatorial mutation of CRR-B and CRR-C (strains *hapX^A/B/C/D^*, *hapX^A/B/C^*, *hapX^B/C/D^*, *hapX^B/C^*) locks HapX in the iron limitation state, i.e. causes a transcriptional iron starvation response in +Fe. Total RNA was isolated from liquid cultures grown in +Fe (A) or +Fe and -Fe (B) at 37 °C. Strains were grown in +Fe without xylose for 17 h followed by 1 h incubation with 0.1% xylose for induction of *hapX* expression (A) and *wt* was grown for 18 h without xylose supplementation (B). Northern blot analyzed genes: *hapX*, iron limitation-induced *mirB* as well as iron limitation-repressed *sreA* and *hemA*. Ethidium bromide-stained rRNA is shown for loading and quality contol.

### C-terminal HapX truncation partially rescues the growth defect caused by combinatorial mutation of the four CRRs

The C-terminal 27 amino acid residues of HapX (amino acid residues 465-491) have previously been shown to be crucial adaptation to -Fe but not hFe ^14^. To investigate a possible link between the HapX C-terminus and the CRRs, we generated mutants expressing *PxylP*-driven *hapX* lacking the C-terminal 27 amino acid residues without (strain *hapX*^464^) or with combinatorial mutation of the four CRRs (*hapX^A/B/C/D;^*^464^). As previously shown with native promoter control ^14^, the C-terminal HapX truncation (strain *hapX*^464^) had no significant effect on growth in hFe (Figure 1). Remarkably, the C-terminal truncation partially rescued the growth defect caused by mutation of the four CRRs with xylose induction in -Fe and +Fe but not in hFe (Figure 1).

Northern blot analysis of mycelia grown in +Fe with xylose induction for 1 h revealed that truncation of the C-terminal 27 amino acid region (strain *hapX^A/B/C/D;^*^464^) attenuated the transcriptional iron starvation response caused by combinatorial mutation of the four CRRs (strain *hapX^A/B/C/D^*), i.e. it decreased the *mirB* transcript level and increased the level of the larger *sreA* transcript, while the *hemA* transcript level was not affected (Figure 5).

**Figure 5:**
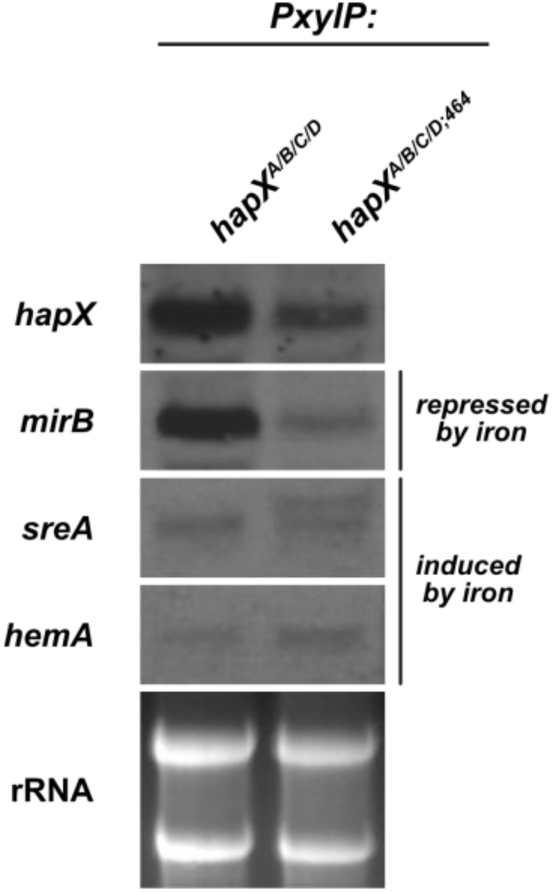
C-terminal HapX truncation (strain *hapX^A/B/C/D,^*^464^) diminishes the transcriptional iron starvation response in +Fe caused by combinatorial mutation of the four CRRs (strain *hapX^A/B/C/D^*). Total RNA was isolated from liquid cultures grown in +Fe at 37 °C for 17 h without xylose followed by 1 h incubation with 0.1% xylose for induction of *PxylP*-driven *hapX* alleles. Northern blot analyzed genes: *hapX*, iron limitation-induced *mirB* as well as iron limitation-repressed *sreA* and *hemA*. Ethidium bromide-stained rRNA is shown for loading and quality contol.

Taken together, the fact that the truncation of the C-terminal 27 amino acid residues, which have previously been shown to be crucial for -Fe adaptation ^14^, partially rescued the growth defect and attenuated the transcriptional -Fe response caused by combinatorial mutation of the four CRRs underlines that the growth defect is caused by the severe -Fe response.

### Expression of *hapX* alleles with combinatorial mutation of CRR-B and -C mutation increases cellular iron accumulation

The transcriptomic analysis indicated that combinatorial mutation of CRR-B and -C causes upregulation of genes involved in iron uptake (Figures 2, 3, 4). To investigate intracellular iron accumulation, the mutant strains expressing the different *hapX* alleles were grown in +Fe liquid culture for 16 h without xylose induction, followed by another 6 h in the presence of xylose to induce expression of the different *PxylP*-driven *hapX* alleles. The schematic experimental setup is shown in Figure 6A. Optical inspection confirmed comparable biomass formation after 16 h of growth without xylose for all strains, consistent with the growth behavior on solid media (Figure 1). However, after the additional 6 h xylose induction period, all strains with combinatorial mutation of CRR-B and -C (*hapX^A/B/C/D^*, *hapX^A/B/C^*, *hapX^B/C/D^* and *hapX^B/C^*; *hapX^A/B/C/D;^*^464^), as well as *hapX^A/C/D^* and *hapX^C^* strains, showed significantly reduced biomass formation compared to *hapX^A/B/D^*, *hapX^A/B^*, *hapX^B^* and *hapX*^464^ and *hapX* (Figure 6B). These results further highlight the importance of HapX CRR-B and CRR-C for the fitness of *A. fumigatus* as seen in the plate growth assays (Figure 1). Among the strains with combinatorial mutation of CRR-B and -C, strain *hapX^A/B/C/D^* showed the most severe growth defect followed by *hapX^B/C^, hapX^A/B/C^* and *hapX^B/C/D^*. The statistically significant difference between *hapX^A/B/C/D^* and *hapX^A/B/C^* (Figure 6B) indicates a function of CRR-D, which was not seen in the growth analyses described above (Figure 1). The *hapX^A/B/C/D;^*^464^ strain displayed significantly increased biomass formation compared to the *hapX^A/B/C/D^* strain (Figure 6B), which is in agreement with the rescue of the growth inhibitory effect seen in the plate growth assays (Figure 1).

**Figure 6:**
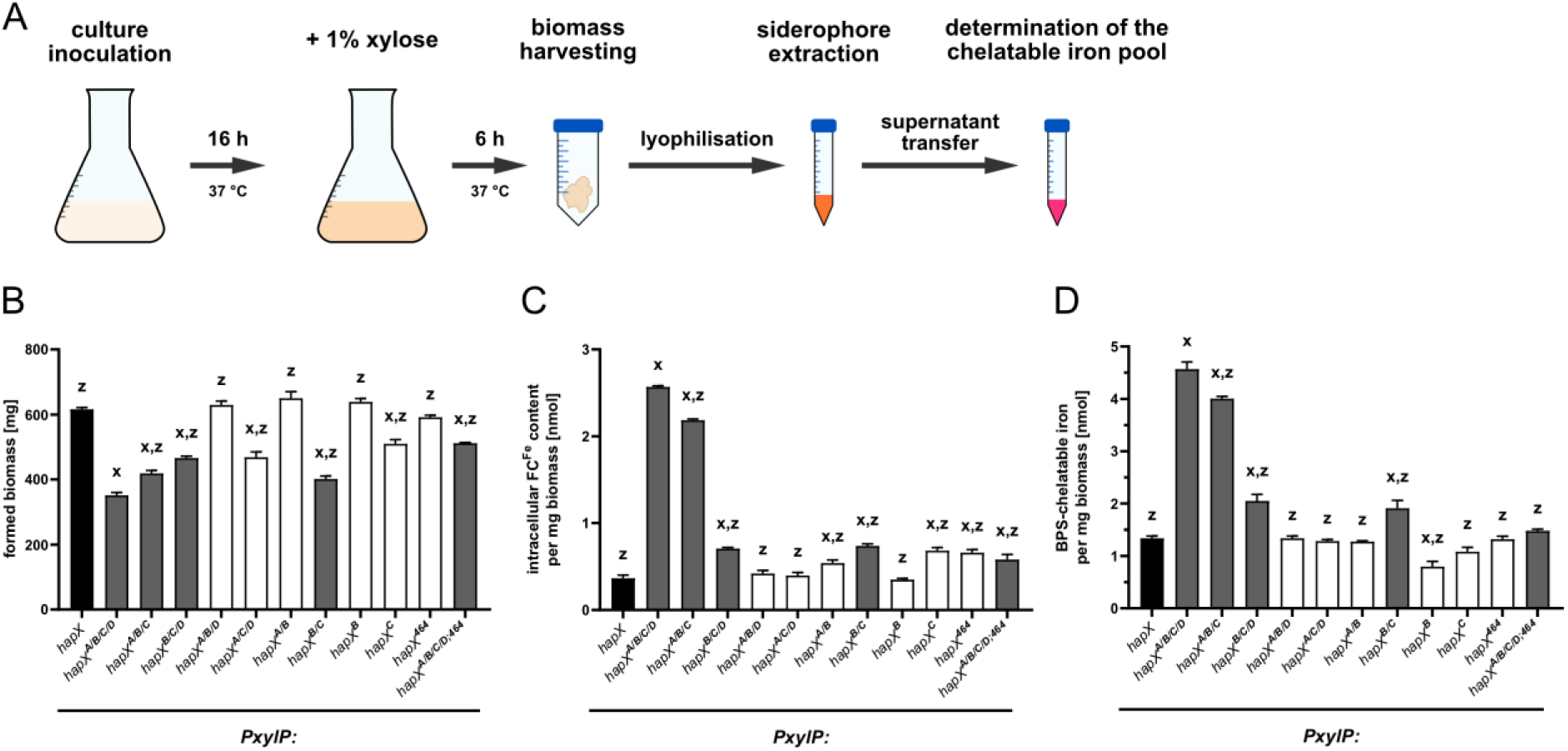
Combinatorial mutation of CRR-B and -C (strains *hapX^A/B/C/D^*, *hapX^A/B/C^*, *hapX^B/C/D^*, *hapX^B/C^*) reduces biomass formation in liquid culture upon xylose supplementation and increases both cellular ferricrocin-complexed iron and the chelatable iron pool. The schematic experimental setup is shown in (A). Liquid cultures were inoculated with the respective fungal strains and grown for 16 h at 37 °C in +Fe conditions, followed by supplementation with 1.0% xylose for *PxylP* induction and continued incubation for additional 6 h. After 22 h of incubation, the harvested biomass was used for estimation of biomass formation (B) and subsequently for the extraction of ferricrocin-complexed iron (C) and the determination of the chelatable iron pool (D). Values shown represent the mean ± standard deviation of biological triplicates. Significant differences, calculated by a two-way analysis of variance (ANOVA), are indicated by “x” for the reference strain *hapX* and “z” for the reference strain *PxylP:hap^XA/B/C/D^*with p < 0.0001.

Further analysis revealed that combinatorial mutation of CRR-B and -C increases cellular levels of iron complexed by the intracellular siderophore ferricrocin, a previously shown indicator of cellular iron accumulation ^15,54,55^: ∼ 7-fold in *hapX^A/B/C/D^*, ∼ 6-fold in *hapX^A/B/C^* and ∼ 2-fold in *hapX^B/C/D^* and *hapX^B/C^* compared to the *hapX* strain (Figure 6C). The statistically significant difference between *hapX^A/B/C/D^*and *hapX^A/B/C^* again indicates a function of CRR-D. C-terminal truncation of HapX (strain *hapX^A/B/C/D;^*^464^) significantly reduced the accumulation of ferricrocin-complexed iron caused by combinatorial mutation of the four CRRs (strain *hapX^A/B/C/D^*). The individual mutation of CRR-C (*hapX^C^*), combinatorial mutation of CRR-A and -B (*hapX^A/B^*) and C-terminal truncation (*hapX*^464^) also resulted in slightly increased intracellular accumulation of ferricrocin-complexed iron (Figure 6C).

Another indicator of cellular iron overload is the chelatable iron pool, i.e. cellular iron not complexed by siderophores, which can be measured photometrically by reddish coloration due to chelation by a chelator such as BPS after reduction by ascorbic acid to ferrous iron ^24^. Only strains expressing *hapX* alleles with combinatorial mutation of CRR-B and -C showed significantly increased chelatable iron pools with *hapX^A/B/C/D^*> *hapX^A/B/C^* > *hapX^B/C/D^* = *hapX^B/C^*(Figure 6D). Again, the statistically significant difference between *hapX^A/B/C/D^* and *hapX^A/B/C^* indicates a function of CRR-D and C-terminal truncation (strain *hapX^A/B/C/D;^*^464^) significantly decreased the chelatable iron pool caused by combinatorial mutation of the four CRRs (strain *hapX^A/B/C/D^*). Remarkably, individual mutation of CRR-B reduced the chelatable cellular iron pool (Figure 6D).

Taken together, the increased levels in ferricrocin-complexed iron and chelatable iron in strains expressing *hapX* alleles with combinatorial mutation of CRR-B and -C are consistent with increased high-affinity iron uptake, as suggested by the gene expression analyses (Figures 2, 3, 4).

### The C-terminus of HapX contains a degron

To analyze the consequences of the described mutations and C-terminal truncation on HapX protein abundance, we performed Western blot analysis in +Fe without xylose induction to allow growth of all mutant strains followed by a 1 h incubation in the presence of 0.1% xylose to induce expression of *PxylP*-driven *hapX* alleles (Figure 7). As reported previously ^10,21^, HapX was barely detectable in *wt* under +Fe conditions and was absent in the *ΔhapX* strain, while *PxylP* control increased the HapX protein level (Figure 7). Compared to the *wt*-like HapX (HapX), the different CRR mutations resulted in similar or slightly higher HapX protein levels, with the combinatorial mutation of CRR-A, -C and D (HapX^A/C/D^) resulting in the highest increase, approximately 6-fold, indicating increased stability of this HapX version as expression of all versions was driven by the same promoter (Figure 7). Remarkably, the truncation of the C-terminal 27 amino acid region, with (HapX^A/B/C/D;464^) or without (HapX^464^) combinatorial mutation of all four CRRs, increased the HapX protein levels approximately 10-fold (Figure 7), suggesting that the truncated region may contain a degron, a protein domain regulating the proteińs half-life by mediating its degradation ^56^. In agreement, fusion of the C-terminal 27 amino acid region of HapX to *PxylP*-driven Venus (strain *Venus^C-termHapX^*) significantly reduced Venus protein levels (Venus *Mr* = 26.8 kDa; Venus^C-termHapX^ *Mr* = 30.3 kDa) in both -Fe and +Fe, as judged by both Venus-derived fluorescence of fungal colonies (Figure 8A) as well as the Venus protein levels in Western blot analysis (Figure 8B), but did not affect the transcript level (Figure 8C). Moreover, Western blot analysis clearly demonstrated degradation of the Venus-HapX fusion protein with the major degradation product having a molecular mass similar to that of Venus. Notably, Venus, like other green fluorescent protein relatives, forms an exceptionally stable 11-stranded β-barrel, which is difficult to degrade leading to quite stable intermediates that are still fluorescent ^57^. Therefore, the Venus protein might be a more stable degradation intermediate.

**Figure 7:**
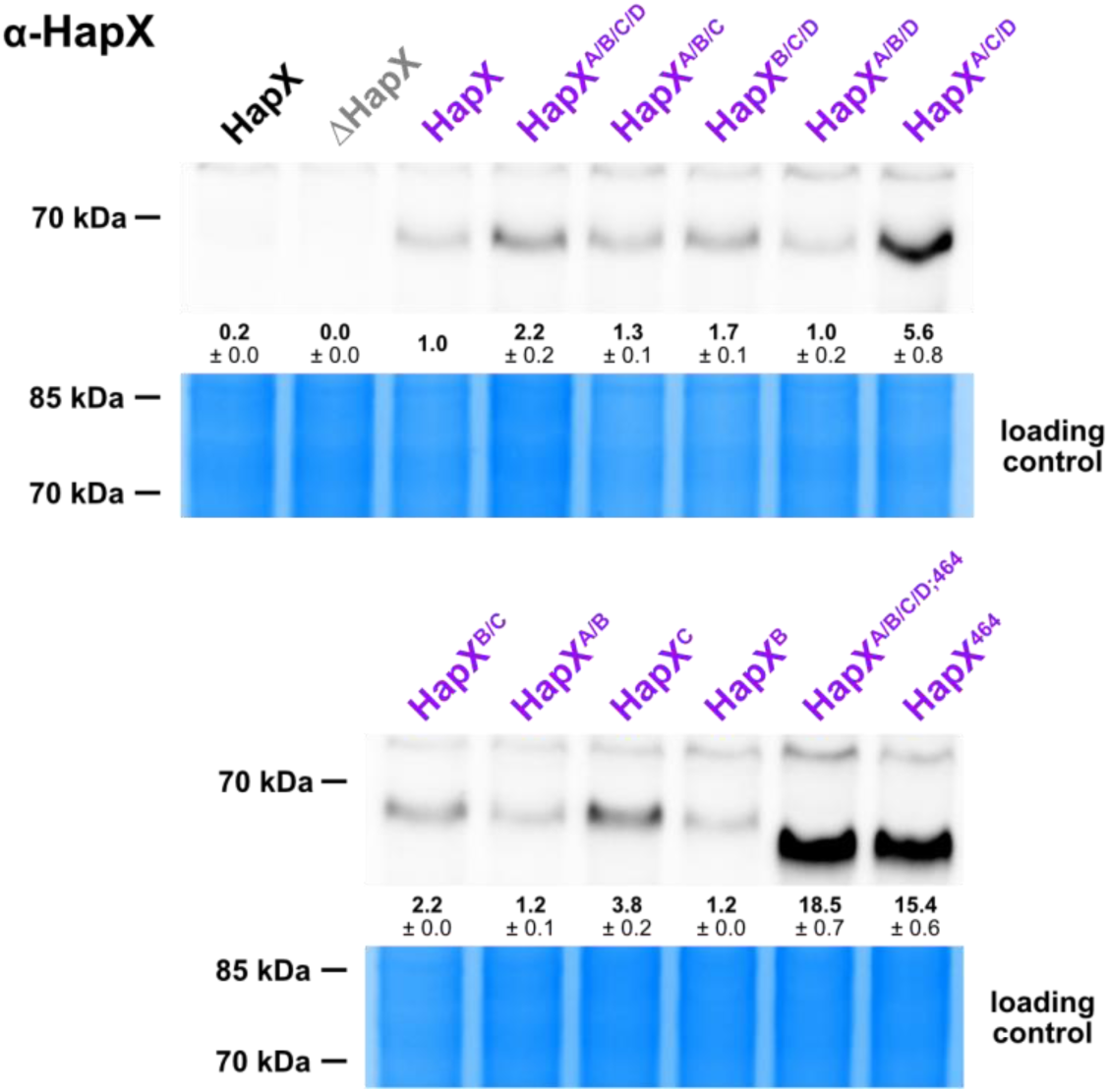
Several mutations of CRRs and in particular C-terminal truncation increases the HapX protein level. Strains were grown in +Fe without xylose for 17 h followed by 1 h incubation with 0.1% xylose for induction of *hapX* expression (same mycelia as used for Northern blot analysis shown in Figure 4) and subjected to Western blot analysis using an α-HapX antibody. All strains with *PxylP*-driven *hapX* alleles are shown in purple; HapX lacking *ΔhapX* in grey serves as the the negative control proving antibody specificity. Production of HapX from the native promoter (*wt* strain) is shown in black. Densitometrically estimated protein amounts were normalised to *PxylP*-driven *wt*-like HapX; numbers represent the mean ± standard deviation. Coomassie stained SDS-polyacrylamide gels are shown as a loading controls.

**Figure 8:**
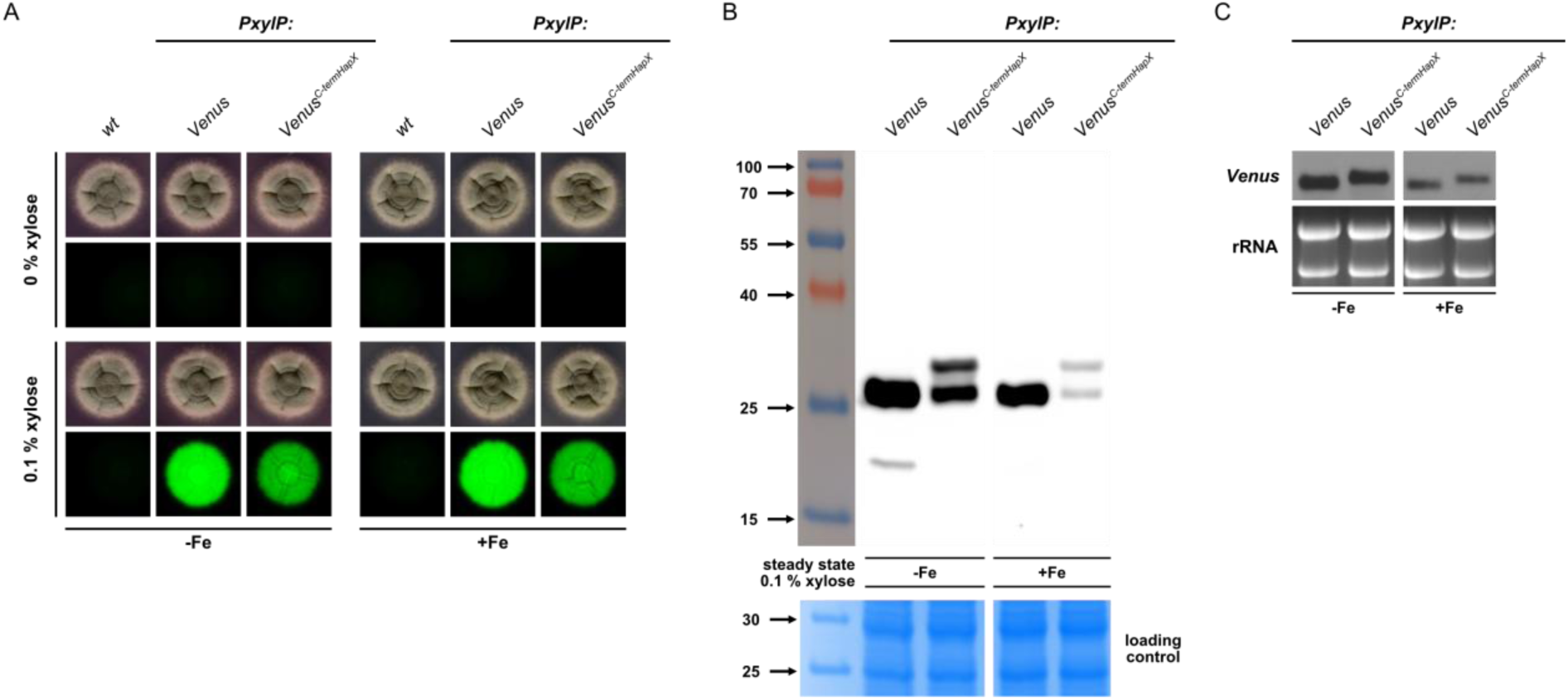
The HapX C-terminus contains a degron. C-terminal fusion of Venus with the C-terminal 27 amino acid region of HapX (strain *Venus^C-termHapX^*) decreases Venus-derived fluorescence intensity of fungal collonies (A) and protein stability in Western blot analysis (B), but does not affect transcript stability (C). Solid (A) and liquid (B) media contained 0.1% xylose for induction of *PxylP*-driven *Venus* alleles. On solid media, 10^4^ conidia were spot-inoculated on medium reflecting iron starvation (-Fe; 200 µM BPS) or iron sufficiency (+Fe; 30 µM FeSO_4_) and incubated at 37 °C for 48 h (A). For Western blot analysis (B) using an α-GFP antibody (Venus *M_r_* = 26.8 kDa; Venus^C-termHapX^ *M_r_* = 30.3 kDa), fungal strains were grown for 17 h in -Fe and +Fe liquid culture; -Fe did not contain BPS. Coomassie stained SDS-polyacrylamide gels are shown as a loading controls. Northern blot analysis was performed with a Venus-specific probe. Ethidium bromide-stained rRNA is shown for loading and quality contol.

## DISCUSSION

HapX is a key regulator of iron homeostasis in *A. fumigatus* that is conserved in most filamentous ascomycetes. It is required for cellular adaptation to both -Fe and hFe, whereas it is dispensable for growth in +Fe. Thus, HapX appears to be able to switch between three functional states to mediate adaptation to -Fe, +Fe (neutral/inactive) and hFe. Its role during -Fe makes HapX important for the virulence of *A. fumi*gatus in mammalian hosts and *Fusarium oxysporum* in plant hosts ^10,58^.

HapX contains four CRRs with four cysteine residues that differ in spacing and amino acid composition and are phylogenetically highly conserved ^4,23^. Previous studies have shown that iron regulation depends on mitochondrial [2Fe-2S] biosynthesis ^24^, that the C-terminal domain of HapX (amino acids 161–491) containing the four CRRs, recombinantly expressed in *E. coli*, exhibits a UV-vis spectrum indicative of the presence of [2Fe-2S] clusters ^17^, that the four CRRs of HapX can complex [2Fe-2S] clusters with different stabilities *in vitro* ^23^, and that the [2Fe-2S] cluster-transferring monothiol glutaredoxin GrxD physically interacts with HapX and appears to mediate removal of [2Fe-2S] clusters from HapX to switch it to the -Fe state ^17^. Taken together, these data suggest that HapX senses the cellular iron state by sensing the abundance of cellular [2Fe-2S] clusters. Consistent with this, mutation of mainly CRR-B, but to a lesser extent also CRR-A and even less CRR-C impaired hFe adaptation of *A. fumigatus*, while mutation of CRR-D had no effect ^14^. These results underline a role for three CRRs in the sensing of hFe by HapX. However, how HapX senses -Fe remained elusive, as individual mutation of any of the four CRRs did not affect -Fe adaptation of *A. fumigatus* ^14^.

Here, systematic combinatorial mutation of the HapX CRRs using conditional expression of the respective alleles with the xylose-inducible *xylP* promoter ^32,33^ revealed that expression of *hapX* alleles with combinatorial mutation of CRR-B and CRR-C (*hapX^A/B/C/D^*, *hapX^A/B/C^*, *hapX^B/C/D^*, *hapX^B/C^*), but not of other combinatorial CRR mutations (*hapX^A/B/D^*, *hapX^A/C/D^*, *hapX^A/B^*), nor individual mutation of CRR-B (*hapX^B^*) or CRR-C (*hapX^C^*), blocks growth regardless of iron availability, i.e. in -Fe, +Fe and hFe conditions (Figure 1). Transcriptome and individual gene expression analyses (Figures 2, 3, 4) revealed that combinatorial mutation of CRR-B and CRR-C results in an transcriptional -Fe response in +Fe, i.e. the induction of genes involved in siderophore-mediated iron acquisition ^15,17^ and the repression of all 33 previously identified HapX target genes involved in iron-consuming pathways ^12^. Taken together, these results suggest that combinatorial mutation of CRR-B and CRR-C locks HapX in the -Fe state. In other words, at least CRR-B and CRR-C, and possibly also CRR-A and -D, lack [2Fe-2S] in -Fe. Consequently, [2Fe-2S] cluster occupancy of CRR-B and/or -C switches HapX out of the -Fe state. Remarkably, of all four CRRs, CRR-B coordinates a [2Fe-2S] cluster with particularly high stability, by far higher than the other three CRRs ^23^. Therefore, CRR-B may be the first CRR sensing iron availability by binding a [2Fe-2S] cluster followed by CRR-C. As suggested by the growth defects in hFe caused by individual mutation of CRR-A, -B, and -C with endogenous promoter control ^14^ and with conditional expression (Figure 1), the switching of HapX to the hFe state then requires the complexation of a [2Fe-2S] clusters also by CRR-A. Notably, in all plate growth assays, mutation of CRR-D was phenotypically inconspicuous in all combinations tested.

In plate assays, all strains expressing *hapX* alleles with combinatorial mutation of at least CRR-B and -C blocked growth in all combinations (Figure 1). However, induction of these *hapX* alleles after uninduced pre-growth in liquid media revealed differences (Figure 6B): Strain *hapX^A/B/C/D^*showed the most severe growth defect followed by *hapX^B/C^, hapX^A/B/C^* and *hapX^B/C/D^*. The statistically significant difference between *hapX^A/B/C/D^*and *hapX^A/B/C^* indicates a function of CRR-D, which was not seen in the growth analyses (Figure 1). Further analysis of these strains revealed that combinatorial mutation of CRR-B and -C increases cellular iron accumulation, i.e. it increased the cellular levels of iron complexed by the intracellular siderophore ferricrocin with *hapX^A/B/C/D^* > *hapX^A/B/C^* > *hapX^B/C/D^* > *hapX^B/C^* (Figure 6C) as well as the chelatable iron pool in a similar order with *hapX^A/B/C/D^* > *hapX^A/B/C^* > *hapX^B/C/D^* ≈ *hapX^B/C^*(Figure 6D). The statistically significant differences in both ferricrocin-complexed iron and the chelatable iron pool between *hapX^A/B/C/D^* and *hapX^A/B/C^*underline a function of CRR-D in iron sensing, which was not seen in the plate growth assays (Figure 1). Remarkably, individual mutation of CRR-B and, to a lesser extent, CRR-C reduced the chelatable cellular iron pool (Figure 6D). Both ferricrocin-complexed iron and the chelatable iron pool are previously shown indicators for iron overload ^15,24,54,55^. The increased cellular iron accumulation in strains expressing *hapX* alleles with combinatorial mutation of CRR-B and -C is consistent with the increased expression of genes involved in siderophore-mediated iron acquisition (Figures 2, 3, 4). Notably, all strains that showed an increased chelatable iron pool (Figure 6D) featured combinatorial mutation of CRR-B and -C and failed to grow on solid media regardless of iron availability (Figure 1), i.e. an increased iron pool was the best indicator of a growth defect in this experimental setup. This may be explained by the fact that the chelatable iron pool has a high potential to cause oxidative stress via Haber-Weiss/Fenton chemistry, whereas chelation of iron by siderophores is thought to prevent the Fenton reaction ^7,59^. Similar to strains expressing *hapX* alleles with combinatorial mutation of CRR-B and -C, depletion of the desulfurase Nfs1, which impairs mitochondrial [2Fe-2S] cluster biosynthesis and consequently iron sensing, was shown to result in a transcriptional -Fe response in combination with an increase in both ferricrocin-complexed iron and chelatable iron ^24^. These results underscore that combinatorial mutation of CRR-B and -C impairs iron sensing via [2Fe-2S] cluster perception.

Previously, the C-terminal 27 amino acid residues of HapX (amino acid residues 465-491) have been shown to be crucial for adaptation to -Fe but not to hFe ^14^, i.e. their truncation impaired growth, siderophore biosynthesis and repression of iron-consuming pathways in -Fe. Truncation of this C-terminal region (strain *hapX^A/B/C/D;^*^464^ compared to strain *hapX^A/B/C/D^*) partially rescued the growth defect caused by mutation of the four CRRs (Figure 1), attenuated the transcriptional -Fe response (Figure 5), and reduced both ferricrocin-complexed iron (Figure 6C) and the chelatable iron pool (Figure 6D). These results underline that the growth defect caused by combinatorial mutation of the four CRRs is indeed caused by the locked HapX -Fe state resulting in activation of high-affinity iron uptake combined with repressed iron consumption due to HapX-mediated repression of genes involved in iron consuming pathways, including repression of CccA-mediated iron detoxification by vacuolar iron deposition ^13^, leading to cellular iron overload and cellular stress. Consistent with these findings, combined mutation of all CRRs caused transcriptional upregulation of several genes encoding stress-responsive regulatory or structural proteins including the mitogen-activated protein kinase MpkB, the basic leucine zipper (bZIP) transcription factors AtfB and AtfD glutathione biosynthetic GshA, several glutathione transferases (Gto1, GstA, GstB), Cu/Zn-superoxide dismutase SodA and nitrosative stress-detoxifying S-nitrosoglutathione reductase GnoA (Figure 3C). Notably, the genes encoding oxidative stress-detoxifying mycelial catalases Cat1 and Cat2 as well as nitrosative stress-detoxifying flavohemoglobin FhpA were significantly downregulated (Figure 3A). However, these genes have previously been shown to be directly repressed by HapX during -Fe ^12,14^. Consequently, the activation of the -Fe state of HapX in +Fe also impairs defense against oxidative and nitrosative stress. The transcriptome analysis also revealed the downregulation of numerous genes involved in ribosome biogenesis and translation (Supplementary Figure S2 and Supplementary Table S1C), and upregulation of five genes involved in autophagy (Figure 3D). This pattern is consistent with repression of TOR complex 1 (TORC; target of rapamycin), a nutrient-sensitive, central regulator of cell growth. Starvation, stress and the drug rapamycin repress TORC1, resulting in repression of protein synthesis, induction of autophagy, exit from the cell cycle, and entry into a quiescent G0 state ^43,44^. Therefore, repression of TORC1 may also be involved in growth inhibition by combinatorial mutation of at least CRR-B and -C.

Western blot analysis revealed that truncation of the C-terminal 27 amino acid residues, with or without combinatorial mutation of all four CRRs, increased the HapX protein levels approximately 10-fold (Figure 7), suggesting that the truncated region may contain a degron, a protein domain regulating the protein’s half-life by mediating its degradation ^56^. In agreement, C-terminal tagging of Venus with the C-terminal 27 amino acid region of HapX significantly reduced Venus protein levels in both -Fe and +Fe, as judged by both Venus-derived fluorescence of fungal colonies (Figure 8A) and Venus protein levels (Figure 8B). Notably, Venus like other green fluorescent protein relatives forms an exceptionally stable 11-stranded β-barrel, which is difficult to degrade leading to quite stable intermediates that are still fluorescent ^57^. Previously, the HapX protein was shown to have a very short half-life of about 15 minutes in -Fe, to be post-translationally modified by ubiquitination, sumoylation and phosphorylation and to physically interact with the monothiol glutaredoxin GrxD and the F-box protein Fbx22 ^17,60^. This newly identified degron represents a new factor influencing the protein stability of HapX and illustrates the complex post-translational regulation of this transcription factor.

Interestingly, the transcriptome analysis also indicated that combinatorial mutation of all CRRs causes upregulation of several gene clusters for biosynthesis of secondary metabolites, i.e. fumarylalanine, pyomelanin/tyrosine degradation, helvolic acid, fumigermin, meroterpenoid pyripyropene, fumitremorgins, and fumagillin/pseurotin (Figure 3E). These gene clusters may be directly activated by the -Fe state of HapX, by the general -Fe response, or by the induced stress that ultimately blocks growth. Notably, the induction of fumarylalanine and pyomelanin/tyrosine degradation has previously been linked to cellular stress ^46,47^.

Taken together, the present study suggests a model for the switching of HapX between its three functional states by the different propensities of the four CRRs for [2Fe-2S] cluster coordination dependent on iron availability. In a living cell, there are multiple HapX molecules that are unlikely to all be in the same functional state, allowing fine-tuning of regulation through the distribution of the three functional states. Most of the HapX homologs from filamentous ascomycetes comprise several CRRs, with CRR-B being the most highly conserved one, suggesting a similar mechanism to *A. fumigatus* HapX ^14,23^. However, there are transcription factors that share only some domains and thus not all functions with HapX: *S. cerevisiae* Yap5 has only two [2Fe-2S] cluster-coordinating CRRs, one of which is highly similar to HapX-CRR-B (67% identity at the amino acid level) and is required exclusively for hFe resistance. In contrast, *Schizosaccharomyces pombe* Php4, which is involved exclusively in adaptation to -Fe, lacks any classical CRR and contains two cysteine residues for binding of a bridging [2Fe–2S] cluster in cooperation with a monothiol glutaredoxin ^4,61^. This data illustrates the modular toolbox used in iron-regulatory transcription factors and underlines the requirement of multiple CRRs for transcription factors enabling adaptation to both -Fe and hFe conditions.

### AUTHOR CONTRIBUTIONS

S.O.: conceptualization, methodology, investigation, data curation and evaluation, visualization, writing – original draft, writing – review and editing.

M.M.: conceptualization, methodology.

H.H.: conceptualization, funding acquisition, investigation, project administration, resources, supervision, visualization, writing – original draft, writing – review and editing.

### DECLARATION OF AI USE

We have not used AI-assisted technologies in creating this manuscript.

## ACKNOWLEDGEMENTS and FUNDING

This research was funded by the Austrian Science Fund (FWF) [Grant-DOI: 10.55776/DOC82] to H.H. For open access purposes, the author has applied a CC BY public copyright license to any author-accepted manuscript version arising from this submission.

The authors acknowledge support from COST Action (European Cooperation in Science and Technology), FeSImmChemNet, CA21115.

## MATERIAL and METHODS

### GROWTH CONDITIONS

*Aspergillus* minimal medium ^62^ containing 1% (w/v) glucose and 20 mM glutamine as carbon and nitrogen sources, respectively, supplemented with iron-free trace elements was used to grow *A. fumigatus* strains. For phenotyping under different iron availabilities on solid media, media were supplemented with 0.2 mM bathophenanthrolinedisulfonic acid (BPS) without iron addition to limit bioavailability and to inhibit reductive iron assimilation ^63^ for -Fe conditions, 0.03 mM FeSO4 to reflect +Fe, or 10 mM FeSO4 to mimic hFe. For solid media, 1.8% (w/v) agar was added, and 10^4^ spores were point-inoculated. For liquid cultures, 100 mL minimal medium in 500 mL Erlenmeyer flasks was inoculated with 10^8^ spores and shaken at 200 revolutions per minute. BPS was not added in -Fe liquid cultures. For induction of the conditional *xylP* promoter (*PxylP*), xylose was added to a final concentration of 0.1% (w/v) ^32,33^. Cultures were generally incubated at 37 °C.

### MUTANT STRAIN GENERATION

In this study, the non-homologous end joining-lacking (*ΔakuA::loxP*) ATCC46645 derivative AfS77, here referred to as *wt*, was used ^64^. Nucleic acid fragments for generation of plasmids were purified either by column purification or by gel extraction (*Monarch PCR & DNA Cleanup Kit* and *Monarch DNA Gel Extraction Kit*, New England Biolabs). Fragments were assembled using the NEBuilder HiFi DNA Assembly or applying site-directed mutagenesis (New England Biolabs); resulting plasmids were amplified in *E. coli* 5-alpha and purified using the *Monarch Plasmid Miniprep Kit* (New England Biolabs).

For generation of a strain expressing *PxylP*-driven *hapX* (strain *hapX*), the 1 kb *hapX* 5’-non-coding region for *hapX* locus-specific targeting was amplified from *wt* genomic DNA using primers *5’hapX flank_fwd*/*5’hapX flank_rev*, the *hph*-*PxylP* cassette was amplified from genomic DNA of strain *hapX^OE^* ^60^ using primers *hph-xylP cassette_fwd*/*hph-xylP cassette_rev* and the *hapX* gene (Afu5g03920) including its native 1 kb 3’-non-coding region was amplified from *wt* genomic DNA using primers *hapX-3’flank_fwd*/*hapX-3’flank_rev*. These fragments were assembled with the backbone of the commercially available plasmid pJet1.2 (Thermo Fisher Scientific) amplified using primers *pJet1.2 backbone_fwd*/*pJet1.2 backbone_rev* applying the NEBuilder approach, yielding plasmid pSO11. The *hph*-*PxylP* cassette contains the hygromycin resistance cassette ^65^, *hph,* and the xylose-inducible *xylP* promoter, termed *PxylP* ^32^, which allows conditional expression of the *hapX* alleles.

The plasmid for the *hapX* version with combinatorial mutations of all four CRRs was based on plasmid pSO11 and was generated by site-directed mutagenesis. Therefore, a single cysteine residue of each CRR was mutated to an alanine residues (C203A, C277A, C353A, C380A) using mutagenesis primers, i.e. *oAfHapX-A2.f/oAfHapX-A2.r*, *oAfHapX-B1.f/oAfHapX-B1.r*, *oAfHapX-C3.f/oAfHapX-C3.r* and *oAfHapX-D2.f/oAfHapX-D2.r*. The resulting plasmid pSO03, encoding the *hapX* allele designated *hapX^A/B/C/D^*(mutated CRRs are indicated in superscript letters), and plasmid pSO11 were subsequently used as templates to generate plasmids encoding the different combinations of CRR mutations; where appropriate, previous cysteine-to-alanine mutations were reversed to the required combinations of CRR mutations. The construction of the other plasmids is not described separately as they were constructed using the site directed mutagenesis strategy as described for pSO03. C-terminal truncations of *hapX* and *hapX^A/B/C/D^*, resulting in *hapX*^464^ and *hap^XA/B/C/D;^*^464^, were also generated using the site directed mutagenesis strategy using primers *oAfhapX-SO7/oAfhapX-SO8*. For fungal transformation the transformation cassettes were PCR-amplified from respective plasmids using primers *oAfhapX-1*/*oAfhapX-2*. After fungal transformation ^66^, transformants were selected using 100 µg/mL hygromycin. For the generation of strains for *PxylP*-driven expression of Venus and a fusion protein of Venus and the C-terminal part of HapX (Venus^C-termHapX^), plasmid pSO52 was generated. Therefore, the following fragments were PCR amplified and assembled using the NEBuilder HiFi DNA Assembly (New England Biolabs): (i) the plasmid backbone containing the *fcyB* 5’- and 3’-non-coding regions, the *PxylP* and the *trpC* terminator derived from plasmid pSO16 ^67^ using primers *fcyB-bb_trpC_fwd/fcyB-bb_xylP_rev*; (ii) the Venus coding sequence derived from plasmid pSO16 ^67^ with the addition of a short linker sequence using primers *Venus_fwd/ Venus_linker-seq_rev*; and (iii) the C-terminal *hapX* coding region from *wt* gDNA using primers *linker-seq_hapX_fwd/hapX_rev*. Plasmid, pSO52 was then used as a PCR template to generate plasmids pSO53 and pSO54. Plasmid pSO53, encoding *Venus* only, was generated by site-directed mutagenesis using primers *pSO53_fwd/pSO53_rev*, deleting the *hapX* coding sequence. Plasmid pSO54, encoding *Venus* fused with the C-terminal 27 amino acid region of HapX, was generated by site-directed mutagenesis using primers *pSO54_fwd/pSO54_rev*. For fungal transformation, the plasmids were linearized by *Not*I restriction digestion and purified by column purification. In this case, no additional selection marker was required as locus-specific integration at the *fcyB* locus was used ^68^. Therefore, transformants were selected using 10 µg/mL 5-fluorcytosine.

The generated strains, plasmids and primers used strains used in this study are summarized in Supplementary Table S2, S3 and S4, respectively. All generated mutant strains were confirmed by Southern blot analysis; the results of the Southern blot analyses and schematic illustrations of the generated genotypes are shown in Figure S3.

### ISOLATION of RNA and DNA; SOUTHERN and Northern BLOT ANALYSIS

Genomic DNA was isolated by PCI extraction and isopropanol precipitation. For Southern blot analysis, DNA digested with specified restriction enzymes, was separated on a 0.7% (w/v) agarose gel and blotted on a Hybond^TM^-N membrane (Cytiva Amersham^TM^) with NaOH for denaturation.

Total RNA was extracted from freshly harvested biomass using TRI reagent (Sigma-Aldrich) according to the manufacturer’s protocol. For RNA gel electrophoresis, 10 µg of isolated RNA were loaded onto 1.2% (w/v) agarose gels with 1.85% (w/v) formaldehyde. For Northern blot analysis, Hybond^TM^-N+ membranes (Cytiva Amersham^TM^) were used.

Hybridization probes for Southern and Northern blot analyses were generated by PCR amplification using digoxigenin-labelled deoxyribonucleotides (Roche); used primers are listed in Supplementary Table S4 and S5, respectively.

### TRANSCRIPTOM ANALYSIS USING RNA SEQUENCING

Illumina RNA library preparation, miCORE mRNA sequencing and bioinformatic analysis was performed by Microsynth AG. Illumina reads were mapped to the reference genome, *A. fumigatus* Af293 ASM265 (NCBI RefSeq assembly: GCF_000002655.1) downloaded from NCBI database ^69^ using STAR version 2.7.11b ^70^. The gene counts were normalized applying DESeq2 ^71^. Differentially expressed genes were filtered using log2 fold change ≤ -1 or ≥ 1 and adjusted p-value < 0.05 as cut-offs. Visualization of the differential gene expression patterns in heat maps was performed using GraphPad Prism version 10.2.3 (GraphPad Software, www.graphpad.com). For functional categorization of the differentially expressed genes, the web-based tool FungiFun (version 2.2.8 BETA) was used ^72^.

### DETERMINATION OF THE CELLULAR FERRICROCIN CONTENT AND THE CHELATABLE IRON POOL

Cellular contents of iron-loaded ferricrocin were determined as described previously ^24,73^. 80 mg lyophilized mycelium was homogenized and resuspended in 1.8 mL sodium-phosphate buffer [pH 7.5; 50 mM] and incubated for 30 min on ice. Cellular debris was removed by centrifugation and 0.8 mL of the supernatant was transferred to fresh tubes and mixed with 0.2 mL of ROTI^®^ phenol/chloroform/isoamyl alcohol (50:24:1; Roth). After phase separation by centrifugation, 0.1 mL of the organic phase was transferred to fresh tubes and mixed with 0.1 mL water and 0.5 mL diethyl ether. After vigorous mixing and centrifugation, the ferricrocin concentration of the lower aqueous phase was measured at a wavelength of 434 nm using a nano spectrophotometer (Thermo Fisher Scientific).

The chelatable iron pool of was quantified in the ferricrocin-depleted supernatant after ROTI^®^ phenol/chloroform/isoamyl alcohol (50:24:1; Roth) extraction and addition of BPS and ascorbic acid to final concentrations of 1 mM each, as previously described ^24^. BPS-complexed ferrous iron was recorded at a wavelength of 535 nm using a molar extinction coefficient of e = 17,000 L mol^-1^ cm^-1^.

### PROTEIN EXTRACTION and WESTERN BLOT ANALYSIS

Total protein was extracted from lyophilized biomass (Triad freeze dryer, Labconco) by alkaline lysis, in combination with trichloroacetic acid mediated protein precipitation, as reported previously ^74^. Total protein extracts were separated electrophoretically on 12 % (w/v) SDS-polyacrylamide gels and transferred onto nitrocellulose membranes (Amersham^TM^ Protran^TM^ Premium 0.45 µm NC, Cytiva Amersham^TM^) using the Trans-Blot® Turbo™ System (Bio-Rad Laboratories).

HapX proteins were recorded using a rabbit α-HapX antibody ^17^, 1:10,000 diluted, in combination with a secondary, peroxidase-coupled α-rabbit antibody (Sigma-Aldrich), 1:20,000 diluted, using ECL detection (Cytiva Amersham^TM^). Protein amounts were estimated densitometrically using Fiji ImageJ ^75^.

Venus proteins were detected using a mouse α-GFP antibody (1:10,000 diluted; Roche) as the primary antibody in combination with a peroxidase-coupled secondary anti-mouse antibody (1:10,000 diluted; Sigma-Aldrich). The ECL reagent (Cytiva Amersham^TM^) was used for detection.

### BIOINFORMATICS

Required gene and protein sequences were downloaded from FungiDB ^76^ or from the NCBI database ^69^. Plasmid and protein sequences were generated and maintained using the cloud-based platform Benchling (Biology Software, 2023). Sequence alignments of genes and proteins were generated using Geneious Prime (2024.0.5; https://www.geneious.com) ^77^.

### STATISTICAL ANALYSES

Statistical analyses were carried out applying GraphPad Prism version 10.2.3 (GraphPad, www.graphpad.com).

## ABBREVIATIONS

BPS: bathophenanthroline disulfonic acid
CRR: cysteine-rich region
-Fe: iron-limiting condition
+Fe: iron sufficient condition
hFe: iron excess condition
*PxylP*: xylose-inducible *xylP* promoter
*wt*: wild type;

