## Supplementary information for "Cooperative [2Fe-2S] cluster-binding regulates the functional transitions of the *Aspergillus fumigatus* iron regulator HapX for adaptation to iron starvation, sufficiency and excess"

### SUPPLEMENTARY FIGURES

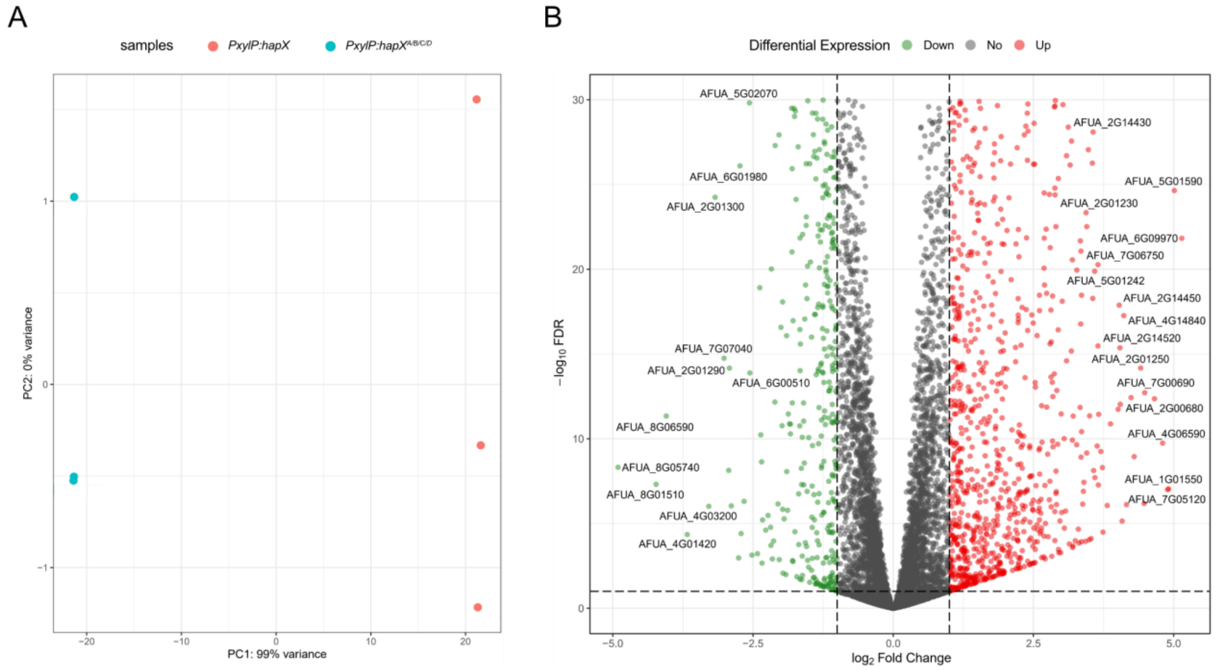


**Figure S1: Global comparison of gene expression in strains *hapX* and *hapX*$^{A/B/C/D}$.** (A) Principal component analysis showing clustering of biological triplicates and differences between the two strains analysed. (B) Volcano blot analysis illustrating the differences in gene expression in *hapX*$^{A/B/C/D}$ compared to *hapX*. Genes downregulated with a log2 fold change ≤ -1 and upregulated with ≥ 1 in *hapX*$^{A/B/C/D}$ compared to *hapX* are highlighted in green and red, respectively. The log2 fold change values are plotted on the x-axis, while the y-axis represents the -log10 false discovery rate (FDR).

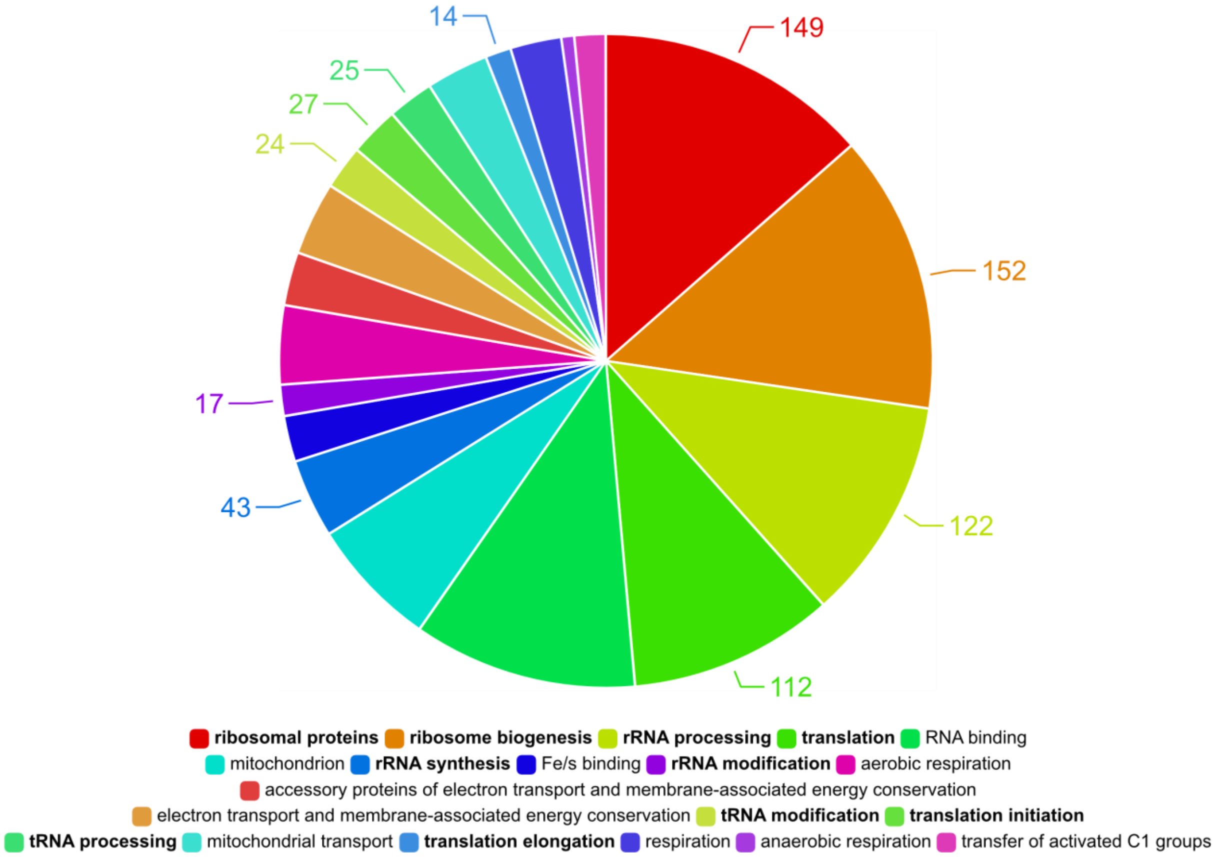


**Figure S2: The majority of downregulated genes in strain *hapX*$^{A/B/C/D}$ are related to ribosome biogenesis and translation.** Functional annotation of significantly downregulated genes in *hapX*$^{A/B/C/D}$ compared to *hapX* was done using FungiFun2 [1]. Functional categories linked to ribosome biogenesis and translation are in bold. Respective genes are listed in Supplementary Table S1C.

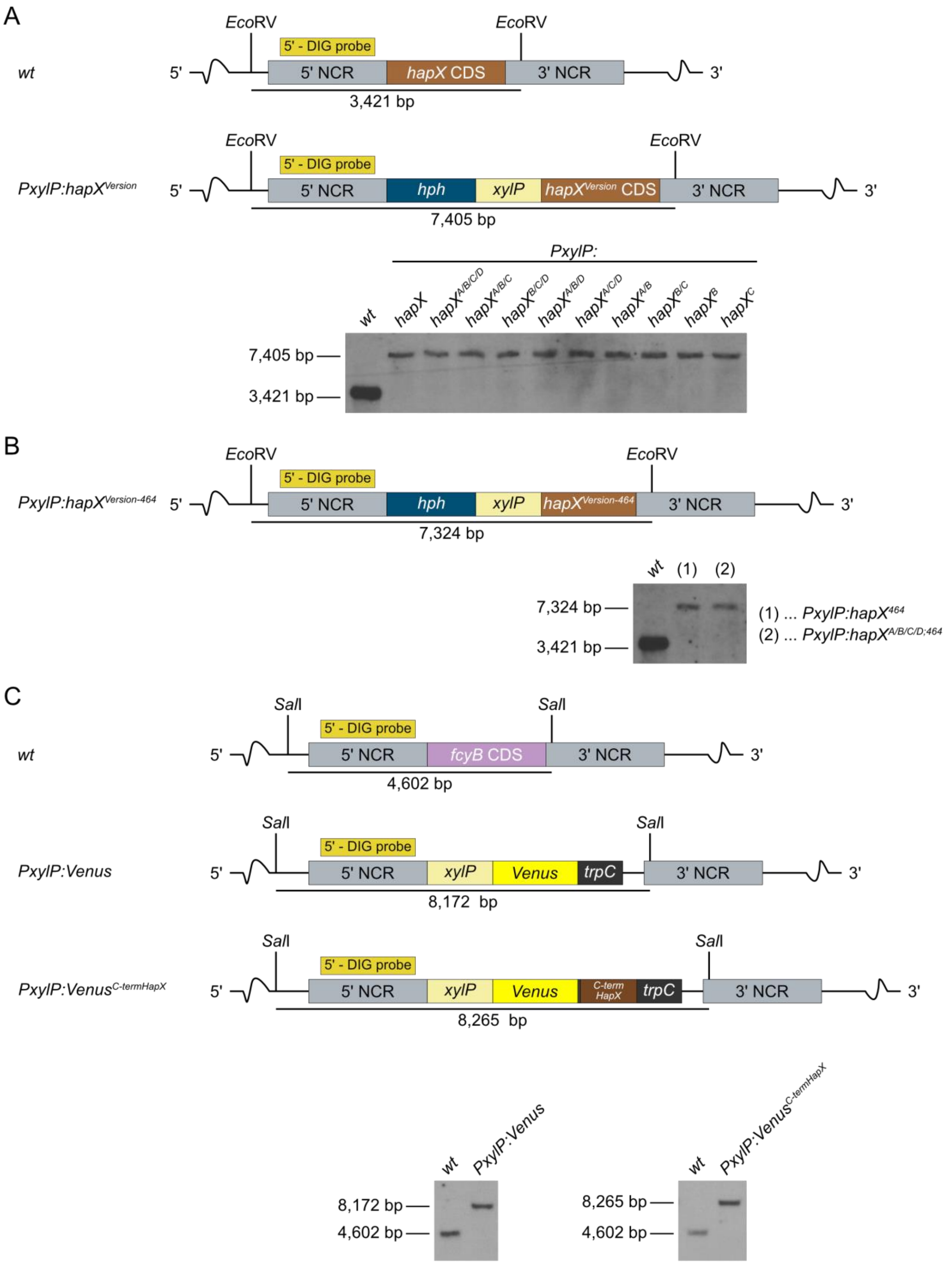


**Figure S3: Schemes of the targeted genomic loci and Southern blot confirmation of the generated mutant strains.** Genomic DNA was digested using specific restriction enzymes and then subjected to Southern blot analysis. (A) *hapX* locus in *wt* and transformed *hapX* CRR mutant strains; digestion using *Eco*RV led to a detected fragment length of 3,421 bp in *wt* and in a 7,405 bp fragment in the mutant strains; (B) in strains encoding C-terminally truncated *hapX* alleles the detected fragment in the mutant strains was slightly smaller, i.e. 7,324 bp. (C) *wt fcyB* locus, used for marker-free integration of the conditionally expressed *Venus* constructs into the *wt* background. *Sal*I digestion resulted in a detected *wt* fragment length of 4,602 bp which increased to 8,172 bp in the *Venus* and to 8,265 bp in the *Venus*$^{C\text{-}termHapX}$ mutant strains, respectively. All Southern blot results are in line with the *in silico* predictions.

### SUPPLEMENTARY TABLES

**Table S1: Transcriptome data of comparative analysis between strains *hapX* and *hapX*$^{A/B/C/D}$.**

***Supplementary Table S1A*** – DESeq2 normalized transcriptome dataset
***Supplementary Table S1B*** – Selected differentially expressed genes in *hapX*$^{A/B/C/D}$ compared to *hapX* in +Fe
***Supplementary Table S1C*** – Functional annotation of significantly downregulated genes using FungiFun2

**Table S2: Strains used in this study.**

| strain | genotype | reference |
|---|---|---|
| *AfS77* (wt) | *ATCC46645, ΔakuA::loxP* | [2] |
| *ΔhapX* | *AfS77, ΔhapX::ptrA* | [3] |
| *PxylP:hapX* | *AfS77, 5'hapX::hph, PxylP:hapX* | this study |
| *PxylP:hapX*$^{A/B/C/D}$ | *AfS77, 5'hapX::hph, PxylP:hapX*$^{A/B/C/D}$ | this study |
| *PxylP:hapX*$^{A/B/C}$ | *AfS77, 5'hapX::hph, PxylP:hapX*$^{A/B/C}$ | this study |
| *PxylP:hapX*$^{B/C/D}$ | *AfS77, 5'hapX::hph, PxylP:hapX*$^{B/C/D}$ | this study |
| *PxylP:hapX*$^{A/B/D}$ | *AfS77, 5'hapX::hph, PxylP:hapX*$^{A/B/D}$ | this study |
| *PxylP:hapX*$^{A/C/D}$ | *AfS77, 5'hapX::hph, PxylP:hapX*$^{A/C/D}$ | this study |
| *PxylP:hapX*$^{B/C}$ | *AfS77, 5'hapX::hph, PxylP:hapX*$^{B/C}$ | this study |
| *PxylP:hapX*$^{A/B}$ | *AfS77, 5'hapX::hph, PxylP:hapX*$^{A/B}$ | this study |
| *PxylP:hapX*$^{C}$ | *AfS77, 5'hapX::hph, PxylP:hapX*$^{C}$ | this study |
| *PxylP:hapX*$^{B}$ | *AfS77, 5'hapX::hph, PxylP:hapX*$^{B}$ | this study |
| *PxylP:hapX*$^{464}$ | *AfS77, 5'hapX::hph, PxylP:hapX*$^{464}$ | this study |
| *PxylP:hapX*$^{A/B/C/D;464}$ | *AfS77, 5'hapX::hph, PxylP:hapX*$^{A/B/C/D;464}$ | this study |
| *PxylP:Venus* | *AfS77, 5'fcyB::PxylP:Venus* | this study |
| *PxylP:Venus*$^{C\text{-}termHapX}$ | *AfS77, 5'fcyB::PxylP:Venus*$^{C\text{-}termHapX}$ | this study |

**Table S3: Summary of generated plasmids in this study and the encoded *hapX* alleles.** Plasmids were either generated applying the NEBuilder or the site-directed mutagenesis (SDM) approach.

| plasmid | encoded *hapX* allele | construction method |
|---|---|---|
| pMMHL79 | *hapX*$^{B}$ | SDM |
| pSO01 | *hapX*$^{A/B}$ | SDM |
| pSO02 | *hapX*$^{A/B/C}$ | SDM |
| pSO03 | *hapX*$^{A/B/C/D}$ | SDM |
| pSO04 | *hapX*$^{A/C/D}$ | SDM |
| pSO05 | *hapX*$^{B/C/D}$ | SDM |
| pSO06 | *hapX*$^{A/B/D}$ | SDM |
| pSO07 | *hapX*$^{A/B/C/D;464}$ | SDM |
| pSO08 | *hapX*$^{B/C}$ | SDM |
| pSO09 | *hapX*$^{464}$ | SDM |
| pSO11 | *hapX* | NEBuilder |
| pSO28 | *hapX*$^{C}$ | SDM |
| pSO52 | *Venus:hapX* | NEBuilder |
| pSO53 | *Venus* | SDM |
| pSO54 | *Venus*$^{C\text{-}termHapX}$ | SDM |

**Table S4: Primers used in this study.** Small letters indicate primer overhangs for plasmid construction or introduction of mutations via site-directed mutagenesis (SDM). PC, plasmid construction; TCA, transformation cassette amplification; SB, Sothern blot probe; SC, sequencing.

| primer | sequence [5' - 3'] | used for |
|---|---|---|
| *pJet1.2 backbone_fwd* | ATCTTTCTAGAAGATCTCCTAC | **PC**<br>➔ **pSO11** [*hapX*] |
| *pJet1.2 backbone_rev* | ATCTTGCTGAAAAACTCG | **PC**<br>➔ **pSO11** [*hapX*] |
| *5'hapX flank_fwd* | ctcgagtttttcagcaagatAGCGACTATAGCCGGATG | **PC**<br>➔ **pSO11** [*hapX*] |
| *5'hapX flank_rev* | taaatggtacGATTACGGATGATGAGAC | **PC**<br>➔ **pSO11** [*hapX*] |
| *hph-xylP cassette_fwd* | atccgtaatcGTACCATTTAATTCTATTTGTGTTTG | **PC**<br>➔ **pSO11** [*hapX*] |
| *hph-xylP cassette_rev* | gtgtagacatGGTTGGTTCTTCGAGTCG | **PC**<br>➔ **pSO11** [*hapX*] |
| *hapX-3'flank_fwd* | agaaccaaccATGTCTACACCTTCAATAGC | **PC**<br>➔ **pSO11** [*hapX*] |
| *hapX-3'flank_rev* | aggagatcttctagaaagatCCTTGGGTCTTGAAGCTTG | **PC**<br>➔ **pSO11** [*hapX*] |
| *oAfHapX-B1.f* | CGTGGATCCCgccGGCTTCTGTTC | **PC – SDM of CRR-B in pSO11**<br>➔ **pMMHL-79** [$hapX^{B}$] |
| *oAfHapX-B1.r* | GCAGGAGACGAAACG | **PC – SDM of CRR-B in pSO11**<br>➔ **pMMHL-79** [$hapX^{B}$] |
| *oAfHapX-A2.f* | CTGCAATGATgcaTCCACATCGCATTG | **PC – SDM of CRR-A in pMMHL79**<br>➔ **pSO01** [$hapX^{A/B}$] |
| *oAfHapX-A2.r* | CCTAACGGTACCTCC | **PC – SDM of CRR-A in pMMHL79**<br>➔ **pSO01** [$hapX^{A/B}$] |
| *oAfHapX-C3.f* | TGCGCGCAGgcgCTTGCAGATCCG | **PC – SDM of CRR-C in pSO01 & pSO11**<br>➔ **pSO02** [$hapX^{A/B/C}$] & **pSO28** [$hapX^{C}$] |
| *oAfHapX-C3.r* | TGTGCCCGGCCCGTTGGC | **PC – SDM of CRR-C in pSO01 & pSO11**<br>➔ **pSO02** [$hapX^{A/B/C}$] & **pSO28** [$hapX^{C}$] |
| *oAfHapX-D2.f* | GTCGGGATGCgcaGGAGGTAAAGGCGC | **PC – SDM of CRR-D in pSO02**<br>➔ **pSO03** [$hapX^{A/B/C/D}$] |
| *oAfHapX-D2.r* | GGGGCAGCGCTAGGG | **PC – SDM of CRR-D in pSO02**<br>➔ **pSO03** [$hapX^{A/B/C/D}$] |
| *oAfhapX*-SO3 | CGTGGATCCCtgtgggTTCTGTTCGGATG | **PC – reverse SDM of CRR-B in pSO03**<br>➔ **pSO04** [$hapX^{A/C/D}$] |
| *oAfhapX*-SO4 | GCAGGAGACGAAACGGAC | **PC – reverse SDM of CRR-B in pSO03**<br>➔ **pSO04** [$hapX^{A/C/D}$] |
| *oAfhapX*-SO1 | CTGCAATGATtgcTCCACATCGCATTGC | **PC – reverse SDM of CRR-A in pSO03 & pSO02**<br>➔ **pSO05** [$hapX^{B/C/D}$] & **pSO08** [$hapX^{B/C}$] |
| *oAfhapX*-SO2 | CCTAACGGTACCTCCTC | **PC – reverse SDM of CRR-A in pSO03 & pSO02**<br>➔ **pSO05** [$hapX^{B/C/D}$] & **pSO08** [$hapX^{B/C}$] |
| *oAfhapX*-SO5 | AgtgtCTTGCAGATCCGCGGAGGAC | **PC – reverse SDM of CRR-C in pSO03**<br>➔ **pSO06** [$hapX^{A/B/D}$] |
| *oAfhapX*-SO6 | GCGCGCATGTGCCCGGCCCG | **PC – reverse SDM of CRR-C in pSO03**<br>➔ **pSO06** [$hapX^{A/B/D}$] |
| *oAfhapX*-SO7 | TGATTTATCGCATCTCTGCTTG | **PC – SDM (C-terminal truncation) in pSO11 & pSO03**<br>➔ **pSO09** [$hapX^{464}$] & **pSO07** [$hapX^{A/B/C/D;464}$] |
| *oAfhapX*-SO8 | CCCACGATCGGTTAAGGG | **PC – SDM (C-terminal truncation) in pSO11 & pSO03**<br>➔ **pSO09** [$hapX^{464}$] & **pSO07** [$hapX^{A/B/C/D;464}$] |
| *fcyB-bb_trpC_fwd* | CCATGGCAGCAGTGATTTC | **PC**<br>➔ **pSO52** [*Venus:hapX*] |
| *fcyB-bb_xylP_rev* | GGTTGGTTCTTCGAGTCG | **PC**<br>➔ **pSO52** [*Venus:hapX*] |
| *Venus_fwd* | atcgactcgaagaaccaaccATGGTCAGCAAGGGCGAG | **PC**<br>➔ **pSO52** [*Venus:hapX*] |
| *Venus_linker-seq_rev* | gtgtagacatGGTTACGGATGACTTGTACAGCTCGTCCATG | **PC**<br>➔ **pSO52** [*Venus:hapX*] |

| | | |
|---|---|---|
| *linker-seq_hapX_fwd* | atccgtaaccATGTCTACACCTTCAATAGC | **PC** ➔ **pSO52** [*Venus:hapX*] |
| *hapX_rev* | tgaaatcactgctgccatggTCATTTGTCGGCAAACCG | **PC** ➔ **pSO52** [*Venus:hapX*] |
| pSO53_fwd | TGACCATGGCAGCAGTGATTTC | **PC – SDM (truncation) in pSO52** ➔ **pSO53** [*Venus*] |
| pSO53_rev | CTTGTACAGCTCGTCCATGC | **PC – SDM (truncation) in pSO52** ➔ **pSO53** [*Venus*] |
| pSO54_fwd | GTCCCAAGGGCCGCTTTG | **PC – SDM (truncation) in pSO52** ➔ **pSO54** [$Venus^{27aaC\text{-}termHapX}$] |
| pSO54_rev | GGTTACGGATGACTTGTACAGCTC | **PC – SDM (truncation) in pSO52** ➔ **pSO54** [$Venus^{27aaC\text{-}termHapX}$] |
| *oAfhapX*-seq1 | TCGGTGGAAAGAAGTGCC | SC |
| *oAfhapX*-seq2 | CGAGTCCGTTTGGGTATC | SC |
| *oAfhapX*-S3 | TTTGTCGGCAAACCGTCG | SC |
| *oAfhapX*-1 | AGCGACTATAGCCGGATG | TCA, SB |
| *oAfhapX*-2 | CCTTGGGTCTTGAAGCTTGCG | TCA |
| *oAfhapX*-4 | ATCAGAGCTGGAGAGGCA | SB |
| *oAfhapX-xylP*.seq | GTATAAGTATCGCCTCCATC | SC |
| *5'fcyB*_fwd | CAGAGAATTGCCAAGCTGGT | SB |
| *5'fcyB*_rev | TAGTTCTGTTACCGAGCCGGCCTGAGTCAATCCCCACCAC | SB |

**Table S5: Primers used for generation of DIG-labelled Northern blot probes.**

| primer | sequence [5' - 3'] | targeted gene | short description |
|---|---|---|---|
| *oAfhapX-seq1* | TCGGTGGAAAGAAGTGCC | *hapX* Afu5g03920 | iron regulatory transcription factor |
| *oAfhapX-S3* | TTTGTCGGCAAACCGTCG | | |
| *oAfmirB1* | AAGCCGAGAAAAAGGGGG | *mirB* Afu3g03640 | TAFC transporter |
| *oAfmirB2* | AACCCAGATGAAGCCCAG | | |
| *oAfsreA5* | CTCAGTACGATCGCTTCC | *sreA* Afu5g11260 | iron regulatory transcription factor |
| *oAfsreA6* | GTTGGACGAGTAGGTAGC | | |
| *oAfhemA-f* | CAAAGGCAAGACTCCACG | *hemA* Afu5g06270 | heme biosynthesis |
| *oAfhemA-r* | GGCATCACCAACCAGAAG | | |
| *Venus_probe_fwd* | GACGTAAACGGCCACAAGTT | *Venus* | yellow-fluorescent protein, Venus |
| *Venus_probe_rev* | GAACTCCAGCAGGACCATGT | | |
